# Comparable theta phase coding dynamics along the transverse axis of CA1

**DOI:** 10.1101/2022.03.16.484582

**Authors:** Aditi Bishnoi, Sachin S. Deshmukh

**Affiliations:** Centre for Neuroscience, Indian Institute of Science, Bangalore; Shiv Nadar Institution of Eminence, Gautam Buddha Nagar

**Keywords:** Hippocampus, Theta, Medial entorhinal cortex, Lateral entorhinal cortex

## Abstract

Topographical projection patterns from the entorhinal cortex to area CA1 of the hippocampus have led to a hypothesis that proximal CA1 (pCA1, closer to CA2) is spatially more selective than distal CA1 (dCA1, closer to the subiculum). While earlier studies have shown evidence supporting this hypothesis, we recently showed that this difference does not hold true under all experimental conditions. In a complex environment with distinct local texture cues on a circular track and global visual cues, pCA1 and dCA1 display comparable spatial selectivity. Correlated with the spatial selectivity differences, the earlier studies also showed differences in theta phase coding dynamics between pCA1 and dCA1 neurons. Here we show that there are no differences in theta phase coding dynamics between neurons in these two regions under the experimental conditions where pCA1 and dCA1 neurons are equally spatially selective. We also show that dCA1 local field potentials (LFPs) show higher power in theta range compared to pCA1 LFPs. These findings challenge the established notion of dCA1 being inherently less spatially selective and theta modulated than pCA1 and suggest that theta-mediated activation of the CA1 sub-networks to represent space is task-dependent.

## Introduction

Oscillations in the theta frequency range (5-10 Hz) are prominent rhythms observed in the rodent hippocampus during REM sleep and exploratory behavior (Vanderwolf, 1969). These oscillations are thought to be essential for the physiological operation of the information processing network involving the hippocampus and the entorhinal cortex. Theta oscillations are also thought to facilitate learning and memory (Colgin, 2013; Nuñez & Buño, 2021). The activity of many neurons in the hippocampus and its anatomically connected regions has been shown to be modulated by theta oscillations (hippocampus: Ranck, 1973; entorhinal cortex: Alonso & García-Austt, 1987; Hafting et al., 2008; head direction cells in the anterior thalamus: Tsanov et al., 2011). Theta oscillations provide temporal windows for coordination of neuronal responses (O’Keefe & Recce, 1993; Skaggs et al., 1996; Jones & Wilson, 2005; Mizuseki et al., 2009) that can enable the efficient transfer of information within large networks and thus support learning and memory.

Spatial representations in the hippocampus and the entorhinal cortex are known to be associated with theta oscillations (Mitchell et al., 1982; Buzsáki & Moser, 2013). Place cells and grid cells exhibit a temporal code for space. As the animal advances through the place or the grid field, the neurons progressively fire at earlier and earlier phases of theta oscillations (O’Keefe & Recce, 1993; Skaggs et al., 1996; Hafting et al., 2008). Certain models have proposed a role for theta oscillations in the generation of spatial code. These models use interference of two independent oscillators with different theta frequencies to generate place fields and grid fields in simulations (O’Keefe & Burgess, 2005; Burgess, 2008; Burgess & O’Keefe, 2011; Barry et al., 2012). As theta coordinates the neural activity of spatially selective cells (Lisman et al., 2005; Maurer et al., 2006; Foster & Wilson, 2007; Wang et al., 2015), studying theta oscillations and theta modulated spiking dynamics can give insights into the network computations underlying neural representations of space.

Proximal CA1 (pCA1; adjacent to area CA2) receives stronger projections from the medial entorhinal cortex (MEC) while distal CA1 (dCA1; adjacent to the subiculum) receives stronger projections from the lateral entorhinal cortex (LEC) (Steward & Scoville, 1976; Witter & Amaral, 2004; Ito & Schuman, 2011; Masurkar et al., 2017). These projections originate from the principal neurons in layer III of MEC and LEC and synapse primarily in the stratum lacunosum-moleculare of CA1 (Steward & Scoville, 1976; Witter & Amaral, 2004; Ito & Schuman, 2011; Masurkar et al., 2017). MEC is believed to send path integration-based spatial information to the hippocampus, while LEC is believed to send sensory derived non-spatial information to the hippocampus (Burwell & Amaral, 1998; Hargreaves et al., 2005). MEC also has stronger theta oscillations in the local field potentials (LFPs) with stronger theta modulation of principal neurons as compared to LEC (Deshmukh et al., 2010). Following these inputs, dCA1 was predicted to be primarily involved in processing the non-spatial aspects of the environment with inherently weaker theta modulated spiking dynamics compared to pCA1, which is considered to be primarily involved in spatial information processing. Some studies have shown these predicted functional differences along the transverse axis of CA1 correlated with the topographical afferent projections from the entorhinal cortex. pCA1 pyramidal neurons have been shown to be more spatially selective and exhibit stronger spike modulation by theta oscillations than dCA1 in simple 1d and 2d environments (Henriksen et al., 2010; Oliva et al., 2016). However, in more complex environments like those with objects, dCA1 can be involved in encoding both spatial and non-spatial information (Burke et al., 2011; Knierim et al., 2014).

Recently, we (Deshmukh, 2021) reported that pCA1 and dCA1 exhibit comparable spatial selectivity on a circular track with distinct textures and multiple visual cues in the periphery. This finding raises 2010; Oliva et al., 2016) may not be universal, and the representation of space along the transverse axis of CA1 may change as a function of the environmental and task conditions. In the current study, we test if theta-mediated neural dynamics continue to show differences between dCA1 and pCA1 in this experimental condition where the difference in spatial selectivity disappears. In contrast to the previous reports, dCA1 exhibits theta phase precession and theta modulation comparable to pCA1. LFP theta power is higher in dCA1, while locomotion speed modulation of theta oscillation is comparable between the two sub-regions. Thus, theta-mediated spatial selectivity in CA1 may be a function of environmental complexity and task demands rather than the entorhinal cortex inputs alone.

## Methods

Neurons recorded along the transverse axis of CA1 from 11 male Long Evans rats were used in this study. Data from these rats have been previously reported (Kumar & Deshmukh, 2020; Deshmukh, 2021). See Deshmukh (2021) for details of the methods described briefly here. All animal procedures were performed in accordance with the National Institutes of Health (NIH) animal use guidelines and were approved by the Johns Hopkins University Animal Care and Use Committee.

### Surgery

Rats were implanted with custom-built hyperdrives over the right hemisphere. These hyperdrives, with 15 independently movable tetrodes and 2 references, had a linear bundle of canulae angled at 35° to target the entire transverse axis of dorsal CA1.

### Behavior and electrophysiology

Animals were trained to run clockwise (CW) on a circular track while the tetrodes were moved daily to position them in the dorsal CA1 pyramidal cell layer. Once the tetrodes were positioned, behavior and neural data were recorded over 5-6 sessions in the following order: STD – MIS – STD – MIS – STD for one rat and STD – STD – MIS – STD – MIS – STD for all other rats (Figure 1A), with 15 laps per session on a circular track (inner diameter: 56 cm, outer diameter: 76 cm). STD stands for the standard configuration of local and global cues, and MIS stands for mismatch configuration where (CCW) by equal amount. The total local-global mismatch angles used for different MIS sessions were 45°, 90°, 135°, and 180°. Each mismatch angle was sampled twice over 4-6 days of recordings. The first STD session of the day was used for all statistical comparisons for STD sessions reported here. The animal’s position in the arena was tracked using LEDs and an overhead camera acquiring frames at 30 Hz. Timestamps where the animal was either running in the opposite direction (CCW) on the track or the animal’s head was detected outside the track were removed.

**Figure 1.**
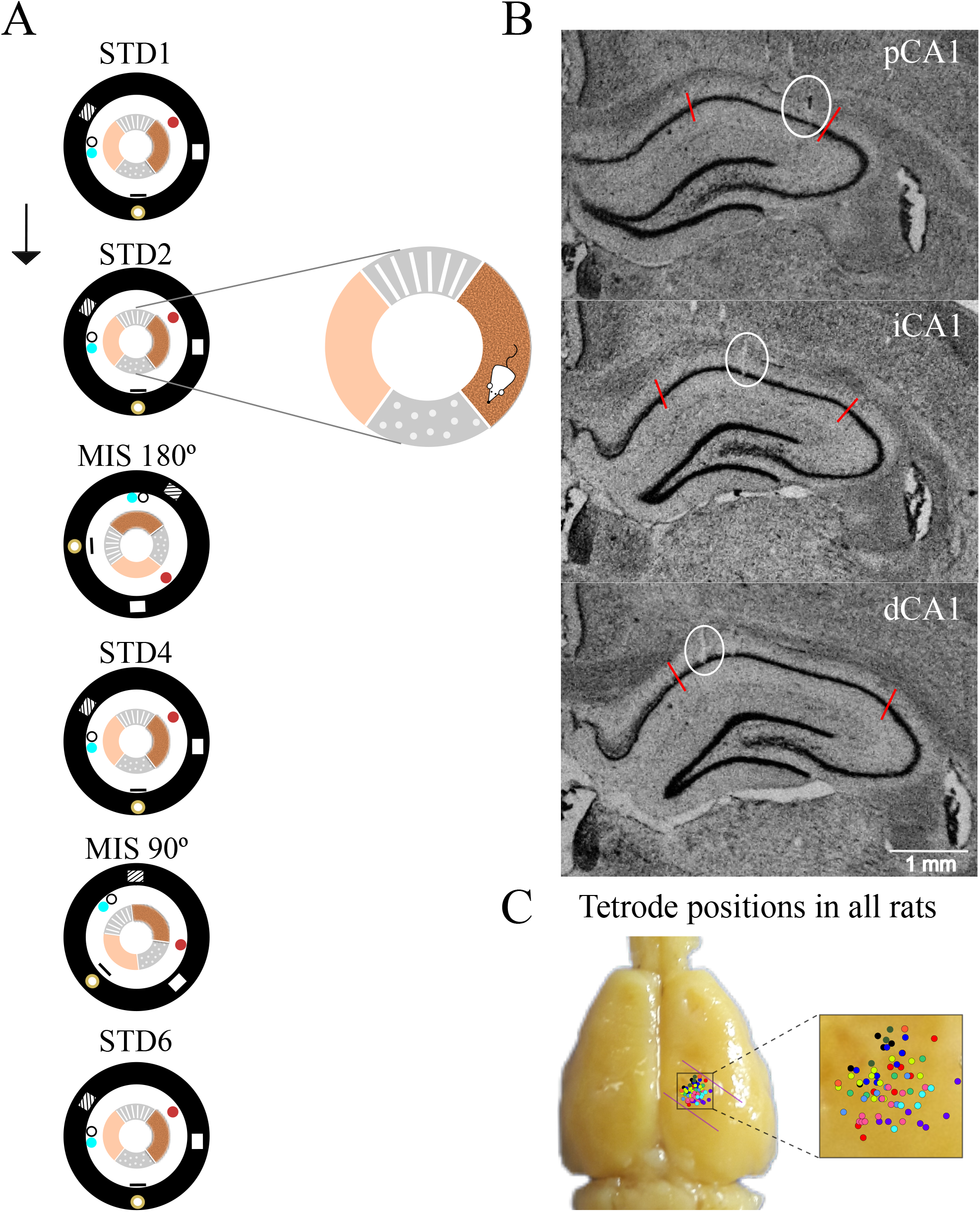
Experimental protocol and recording locations. **(A)** Local and global cue mismatch protocol. Four different textures were used as local cues on the circular track; six global cues were placed along the curtain represented by a black circle around the track, either on the floor or hung from the ceiling. Rats were trained to run CW on the circular track. STD sessions where the local and global cues were maintained in their familiar configuration were interleaved with MIS sessions where the local cues were rotated CCW, and global cues were rotated CW by equal amount. Mismatch angles of 45°, 90°, 135°, or 180° were pseudorandomly selected for a particular MIS session to sample each mismatch angle twice over 4 days of experiments. The arrangement of cues for MIS 180° and MIS 90° is shown here. The arrow marks the order in which the sessions were recorded. **(B)** Coronal sections showing examples of recording locations along the transverse axis of CA1. White circles highlight the example tetrode tracks; red lines mark the CA1 boundaries. Top: an example of a tetrode in pCA1, Middle: iCA1, Bottom: dCA1. **(C)** Recording locations from all rats. Dots of a single color represent tetrodes from a single animal. Adapted from Deshmukh, 2021.

An analog wireless transmitter (Triangle Biosystems International, Durham, NC) was used to collect neural data. Digitization and data processing was done using Cheetah data acquisition system (Neuralynx Inc., Bozeman, MT). The analog signals were amplified 1000-10000 fold, high pass filtered (600-6000 Hz), and digitized at 32000 Hz. Each time the signal crossed a pre-set amplitude threshold, 32 sample points were recorded around the threshold passing event (8 points before the threshold passing event and 24 points after the event at 32 KHz i.e., 1 ms) on all 4 tetrode channels as spike data. For LFP recordings, signals were digitized at 1000 Hz after amplifying 500-2000 fold and bandpass filtering between 1-475 Hz.

A custom manual cluster cutting software (WinClust, J.J. Knierim, Johns Hopkins University) was used to identify single units. Peak, valley, and energy of the spike waveforms on all 4 channels of a tetrode were used to cluster spikes. Putative interneurons with a mean firing rate of ≥ 10 Hz were excluded from further analysis. An isolation quality score was assigned to each unit on a subjective scale of 1 (poorly isolated) to 5 (very well isolated) based on how well the cluster was separated from the neighboring clusters and the background. This score did not use the spatial firing characteristics of the units. Clusters with a quality score of 3 or better were used from all three CA1 sub-regions.

### Place field analysis

To create a linearized firing rate map representing activity of a neuron on the track, the circular track was binned into 360 × 1° bins (1° = 0.57 cm at the mean diameter of the track). The number of spikes a neuron fired while the rat occupied a particular bin was divided by the time the rat spent in that bin to obtain firing rate of that neuron in that bin. This map was smoothed for calculating spatial information score (bits/spike; Skaggs et al., 1996) using adaptive binning. The probability of obtaining the neuron’s spatial information score by chance was determined using a shuffling procedure. For this, a random time lag (minimum 30 s) was added to the neuron’s spike train to created from this shifted spike train to compute spatial information (Skaggs et al., 1996). A chance distribution of spatial information scores for the observed spike train was obtained by repeating this procedure 1000 times. The significance of the observed spatial information was determined from the chance distribution by counting the number of times the trials with random lags had spatial information scores greater than or equal to the observed score. Only putative pyramidal cells with statistically significant (p < 0.01) information scores > 0.5 bits/spike were used in further analysis of theta modulation and theta phase precession in the neuron’s place field. This threshold was empirically determined to select spatially responsive cells with well-defined place fields necessary for this analysis.

Place fields were detected from Gaussian smoothed (window size = 30°, σ = 9°) linearized rate maps. This σ (and the corresponding window size) is larger than the 3° σ used earlier (Deshmukh, 2021). We chose to use this large σ to ensure that the spikes at the end and especially at the beginning of the place field (where the firing rates are very low and rising slowly) are not missed under the threshold used for place field detection. This effect is prominent in larger (> 105°) place fields. Using the earlier 3° σ (Deshmukh, 2021) leads to loss of these spikes but does not substantially alter the results described here. Furthermore, large σ of around 5 cm (∼ 9° on our track) is often used for place field detection (Haggerty & Ji, 2015; Oliva et al., 2016). Place fields were defined as having at least 10 contiguous 1° bins (corresponding to 5.7 cm at the mean diameter of the track) with firing rates ≥ 0.3 Hz and ≥ 10% of the peak firing rate for the field. Each individual place field had to have a peak firing rate ≥ 1 Hz and ≥ 30% of the overall peak firing rate of the neuron, with at least 50 spikes for the place field to be included in the analysis.

### Theta detection and theta phase assignment

To obtain modulation of spiking activity by local theta oscillations, every time a neuron fired, phase of theta oscillations on the same tetrode was assigned to that spike. Peaks were detected on LFP signals filtered in the theta range (5-10 Hz) and assigned a phase of 0/360°. Each spike was assigned a phase between the consecutive peaks by linear interpolation (Skaggs et al., 1996). Epochs with at least 3 contiguous theta cycles, each with a peak amplitude ≥ mean + 0.8σ of the signal amplitude were identified as strong theta events. This criterion was determined by visual inspection of theta epochs to capture the strong theta events across tetrodes across rats. Only spikes that occurred during strong theta events were assigned a theta phase and included in further analysis. Spike theta were highly correlated with phases estimated by Hilbert transform (mean Pearson’s correlation 0.96 ± 0.02 standard error of mean). Using Hilbert transform to determine instantaneous theta phase instead of linear interpolation does not substantially alter the results.

### Theta Phase Precession

Spike phases for two theta cycles were plotted against the position of the animal in the place field to visualize an entire cycle of phase precession. Place fields with overlapping precession cycles were removed manually from further analyses (Supplementary Figure S1). To calculate the strength of precession, the phase vector for each place field was rotated in steps of 1° with respect to the position for the entire phase cycle of 360° (O’Keefe & Recce, 1993). Linear regression between phase and position was performed for each rotation. Out of the 360 linear fits, the regression line with the highest r^2^ (i.e., the highest explained variance of theta phase by the position of the animal in the field) was taken as the best fit for that place field. The corresponding r^2^ value was considered the strength of phase precession for that place field. For performing statistics on all place cells, the place field with the highest r^2^ was chosen for cells with more than one place field. Slope and range of phase precession were also calculated for place fields that showed significant phase-position correlation (i.e., the slope of linear regression fit was significantly non-zero at p < 0.01).

### Modulation of Spiking activity by local theta oscillations

#### Theta Modulation Index (TMI)

Spike phase histograms were generated with a bin size of 36° and smoothed by a Gaussian window (window size = 7 bins, σ = 0.5 bins) for each place field. TMI was defined as the peak to valley difference of the peak normalized histogram. If the spikes are not modulated by theta oscillations, their theta phases are expected to be distributed uniformly. A randomization procedure was employed to estimate the probability that the observed TMI for a place field could be obtained by chance from a uniform distribution. Each spike in the place field was randomly assigned a phase value between 1° to 360° from a uniform distribution to calculate a TMI and the process was repeated for 1000 iterations. The significance of the observed TMI was determined using the proportion of random trials with TMI greater than or equal to the observed TMI. A significance threshold of p < 0.05 was used to identify theta modulated neurons. For performing statistics on all place cells, the place field with the highest TMI was chosen for cells with more than one place field.

#### Phase Precession Adjustment of Theta Modulation Index (TMIa)

To discount the influence of phase precession on TMI calculation, we measured the deviation of the spike phases from the linear phase precession fit. The deviation values could range from −360° to 360°. All negative deviation angles were converted to a 0°-360° scale by adding 360° to the negative values, and a histogram of phase deviations was generated using binning and smoothing as described for TMI. TMIa was taken as the peak to valley difference of the peak normalized phase deviation histogram. For performing statistics on all place cells, the place field with the highest TMIa was chosen for cells with more than one place field. The probability of obtaining this TMIa by chance (significance criterion: p < 0.05) was tested against a chance distribution of TMIa. A randomized TMIa was generated by randomly assigning a phase deviation value between −360° to 360° to each spike. This process was repeated 1000 times to get a chance distribution of TMIa.

### Spectral Analysis

Power Spectra were obtained from LFPs using the MATLAB pspectrum function. To accurately estimate the theta peak parameters from the power spectra, the 1/f^n^ component was estimated and subtracted using an open-source Python toolbox, FOOOF (Donoghue et al., 2020; https://github.com/fooof-tools/fooof, release 0.1.2). The FOOOF algorithm was implemented on the power spectrum of each tetrode, using the ‘knee’ parameter in the module over the frequency range of 3-100 Hz for estimating the aperiodic (1/f^n^) fit. The 1/f^n^ ‘background fit’ returned by the program was subtracted from the original spectrum to obtain a ‘flattened’ spectrum (Supplementary Figure S2). The ‘flattened’ spectra were used to estimate total power in the theta range (5-10 Hz) and frequency with the highest theta power.

### Statistics

Projections from LEC and MEC are expected to overlap in iCA1 (Steward, 1976; Steward & Scoville, 1976). Thus, iCA1 was excluded from all statistical comparisons by prior design. Since we did not have an a priori reason to believe that the data were distributed normally, non-parametric tests were used for statistical comparisons between pCA1 and dCA1: Wilcoxon signed rank test for paired comparisons and Wilcoxon rank sum test for unpaired comparisons. To control for the possibility that the observed lack of statistical difference was due to weak non-parametric tests used here, post-hoc parametric tests (paired t-test for paired comparisons and two sample t-test for unpaired comparisons) were performed for distributions that did not significantly differ from a normal test which is suitable when the parameters of the normal distribution are unknown.

The recordings were performed on every animal over 4-6 days with 5-6 sessions on each day. Tetrodes recording multiple units were left undisturbed from one day to another, while those without units were moved ∼16-32 µm to increase the yield. Units were tracked through the STD and MIS sessions on a recording day, but were not tracked from one recording day to another. This resulted in repeat sampling of a subset of units across sessions and across days. Thus, corrections for multiple comparisons could not be performed, as such corrections assume independence of samples. Instead, inference was drawn from patterns of low p values (p < 0.05) across multiple comparisons. In all analyses where multiple comparisons were performed simultaneously, no conclusions were drawn based on single comparisons.

Effect sizes are reported as difference in the medians (dCA1 subtracted from pCA1 in all cases) with 95% Confidence Interval (CI) calculated using bootstrapping methods in MATLAB. Standardized mean differences (Cohen’s d; Cohen, 1988) are also reported for cases where post-hoc tests revealed samples to be normally distributed.

Frequentist statistics were complemented by Bayesian statistics. Bayes Factors using Wilcoxon test were calculated in JASP (JASP Team, 2022) with default prior distributions. The Bayes factor is a likelihood ratio that compares the evidence for a null hypothesis H_0_ against an alternative hypothesis H_1_ (Schönbrodt & Wagenmakers, 2018). We denote the ratio as BF_10_ (“H_1_ over H_0_”). BF_10_ = 4 indicates that the data are four times more likely under H_1_ than under H_0_ i.e., H_1_ is a better probabilistic prediction for the observed data than H_0_. Thus, BF_10_ > 1 supports H_1_ over H_0_, while BF_10_ < 1 supports H_0_ over H_1_. Evidence in favor of an effect is considered anecdotal if BF_10_ < 3, moderate if 3 < BF_10_ < 10, strong if 10 < BF_10_ < 30, very strong if BF_10_ > 30, and extremely strong if BF_10_ > 100 (Lee & Wagenmakers, 2013).

## Results

LFP and single-unit activity were recorded along the transverse axis of dorsal CA1 of 11 Long Evans rats using independently movable tetrodes. These recordings were performed as the animals ran on a circular track with four distinct textures that served as local cues and six distinct global visual cues along the surrounding circular curtain. STD sessions were interleaved with MIS sessions where the (Figure 1A). We compared theta modulated spiking dynamics between pCA1 and dCA1 neurons during STD and MIS sessions. Locations of the tetrodes along the transverse axis were confirmed using histological reconstruction at the end of the experiment (Figure 1B,C).

### Comparable theta phase precession between pCA1 and dCA1

Hippocampal place cells often display theta phase precession by progressively shifting the theta phase in which the place cells fire as the animal passes through the neurons’ place fields. To study phase precession of a neuron, we first identified theta events throughout a session on every tetrode. LFPs from the tetrode from which a given neuron was recorded were used to measure theta oscillations local to that neuron. The phase of local theta oscillation was estimated for each spike in a neuron’s place field during strong theta events (Figure 2A). In contrast to the previous report from 1D environment (Oliva et al., 2016), we found the strength of phase precession to be statistically indistinguishable between pCA1 and dCA1 cells in our experimental protocol (Figure 2B, C; Wilcoxon rank sum test; z = −0.4, rank sum= 9773, p = 0.688; pCA1 median 0.277, dCA1 median 0.29, median difference = −0.013 with 95% confidence interval= [−0.07, 0.04]). Post-hoc Lilliefors test showed that the samples are normally distributed (p = 0.91 and 0.48 for pCA1 and dCA1, respectively). Post-hoc t-test (t (202,0.12) = −0.27, p = 0.78) and Cohen’s d = −0.037 (confidence interval = [−0.311, 0.236]) confirmed that strength of phase precession is indistinguishable between pCA1 and dCA1. A place cell was considered to have significant phase precession if the phase-position correlation was significant (p < 0.01) in at least one of its place fields. Almost all neurons in CA1 showed significant theta phase precession (94/97 cells in pCA1, 86/91 cells in iCA1 and 101/107 cells in dCA1). The proportions of cells showing significant theta phase precession were statistically indistinguishable in pCA1 and dCA1 (Figure 2D; chi-square test for independence; χ^2^ = 0.76, p = 0.382). For the theta phase precessing neurons, the dynamics of precession (slope and range of precession) also showed no significant difference between the two sub-regions (Figure 2E; Wilcoxon rank sum test, slope: z = 1.3, rank sum = 9726, p = 0.192; pCA1 median −5.36 degrees/cm, dCA1 median −5.71 degrees/cm, median difference 0.35 [−0.32, 1.04] degrees/cm; range: z = −1.31, rank sum = 8694, p = 0.189; pCA1 median 210.8°, dCA1 median 222.1°, median difference −11.3° [−23.9°, 5.6°]). Test for normality failed for post-hoc parametric tests for both slope and range of precession (Lilliefors test p < 0.05, except for slope of precession in dCA1, p = 0.43), hence Cohen’s d was not determined. Absence of a difference in theta phase precession dynamics between pCA1 and dCA1 contrasts previous studies selectivity between the two sub-regions in the same experimental protocol (Deshmukh, 2021).

**Figure 2.**
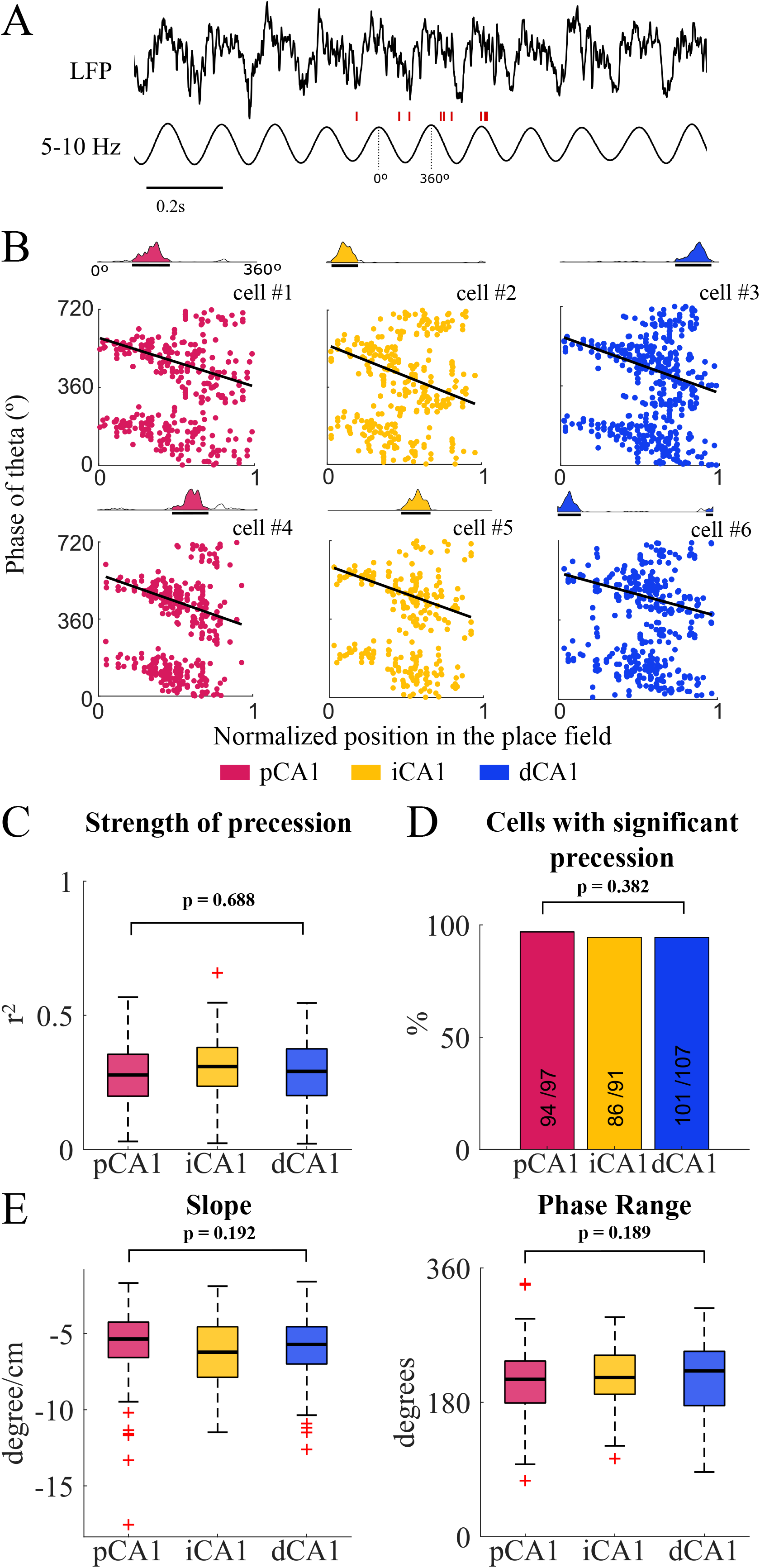
Theta phase precession. **(A)** Example raw LFP trace (top) and corresponding filtered signal (bottom). Vertical red lines represent spikes from one of the neurons that were isolated on the same tetrode. **(B)** Example cells from each region along the CA1 transverse axis showing theta phase precession. For each cell, top trace: firing rate of the cell as a function of position on the linearized circular track. Black lines below the rate map mark the detected place field. Bottom: scatter plot of spike theta phases as a function of the position of the animal in the place field. Position in the place field has been normalized for display purposes. 0 and 1 mark the beginning and end of the place field, respectively. **(C)** Box and whisker plots showing the distribution of precession strength (r^2^, i.e. the variance in spike phase explained by position in the place field) for all place cells recorded in STD session. Medians are represented by horizontal lines in each box. Each box spans from the 25^th^ to the 75^th^ percentile (the difference is the interquartile range-IQR). Outliers, marked with red crosses, are values more than 1.5 times the IQR away from the top or bottom of the box. Whiskers extend above and below each box marking the range between the non-outlier maximum and the non-outlier minimum. **(D)** Proportion of cells with significant theta phase precession in at least one of their place fields. **(E)** Distribution of slope and range of phase precession for the cells showing phase precession.

### Theta modulation along the transverse axis of CA1

A theta modulation index (TMI) was calculated for each place field of a neuron as the difference between the peak and the valley of the spike theta phase histogram normalized by the peak (Figure 3A). We found TMI to be significantly higher in pCA1 than dCA1 (Figure3B; Wilcoxon rank sum test; z = 2.74, rank sum = 11097, p = 0.006; test for normality passed for post-hoc parametric tests, Lilliefors test (p = 0.09 and 0.72 for pCA1 and dCA1, respectively). Post-hoc t-test also showed a significant difference (t (202,0.16) = 2.94, p = 0.004) with small to medium effect size (pCA1 median 0.7, dCA1 median 0.63, median difference of 0.067 [0.002,0.143]. Note that the lower bound of 95% CI is close to 0; post-hoc Cohen’s d= 0.41 [0.13,0.68] is in the small (0.2) to medium (0.5) range (Cohen, 1988, p25). However, the proportion of cells modulated by theta oscillations was statistically indistinguishable between pCA1 and dCA1 (Figure 3C; chi-square test for independence; χ^2^ = 0.25, p = 0.62; proportion of cells with significant TMI in pCA1 = 80/97, iCA1 = 76/91, dCA1 = 91/107). This similar proportion of theta modulated cells in both sub-regions is in contrast with previous reports where pCA1 has been shown to have higher proportion of cells modulated by local (Oliva et al., 2016) as well as MEC (Henriksen et al., 2010) theta oscillations compared to dCA1. Similar proportions of significantly theta modulated cells but lower values of TMI in dCA1 can be explained if cells in dCA1 exhibit phase precession over a wider range of theta phases. The range of phase precession was not significantly different between pCA1 and dCA1 but showed a trend of dCA1 being higher than pCA1 (Figure 2E; median difference skewed towards dCA1, −11.3° [−23.9°, 5.6°]).

**Figure 3.**
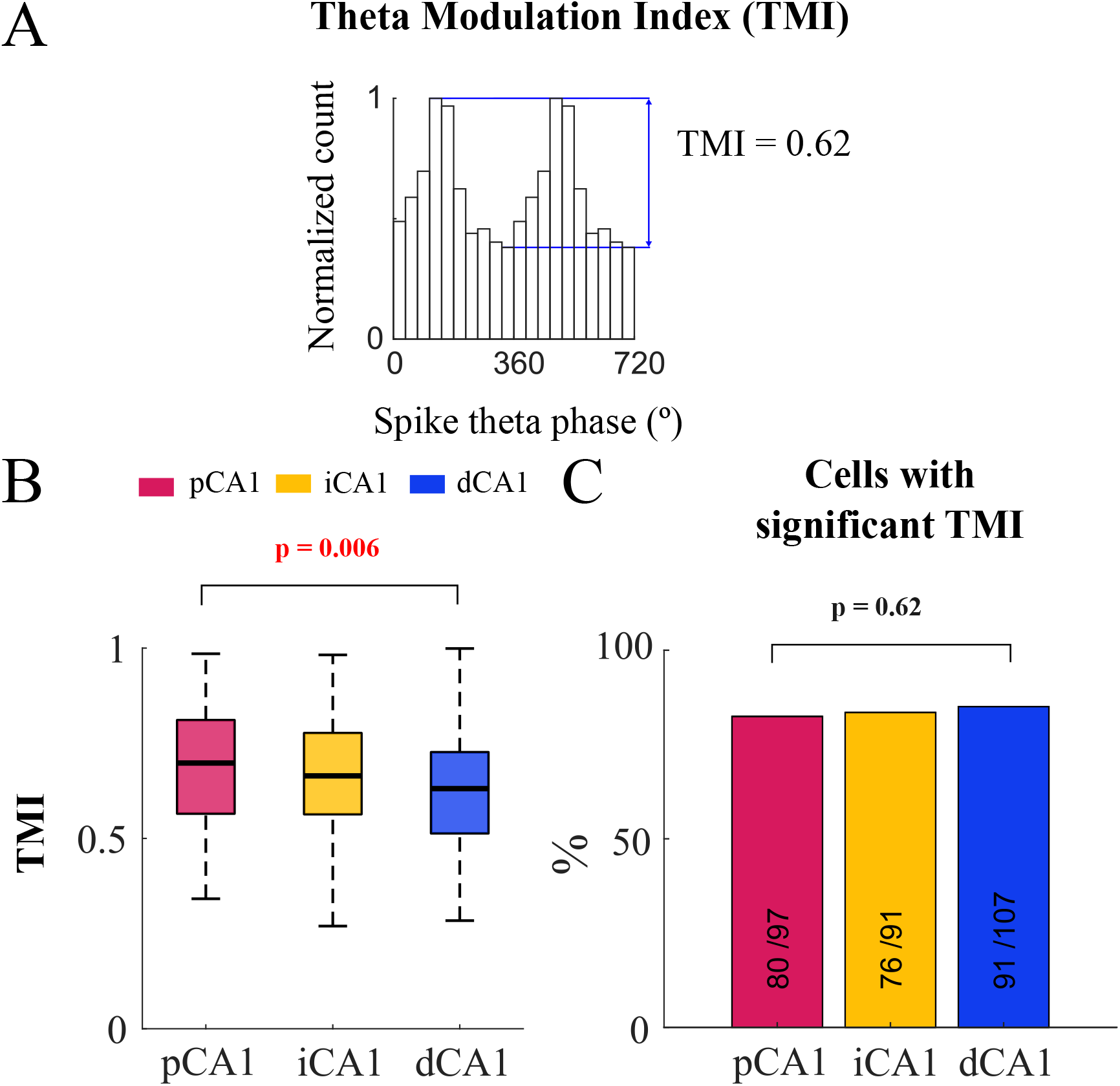
Theta modulation. **(A)** Calculation of Theta Modulation index (TMI) from the spike phase histogram for 2 cycles of theta oscillation for an example neuron. **(B)** Box and whisker plots of the distribution of TMI for all place cells recorded in STD session. **(C)** Proportion of cells with significant TMI in at least one of their place fields were similar in pCA1 and dCA1 (numbers inside bars indicate numbers of cells with significant TMI and the total number of cells).

In theta phase precession, the preferred phase of theta oscillation in which the neuron fires changes with the rat’s position. Thus, the depth of the theta modulation histogram (which averages theta phases in which the neuron fires across positions) gets shallower in precessing neurons (Figure 4A). This weakens our estimate of modulation of the neuron’s activity by theta oscillations using TMI. To compensate for the dependence of neuron’s preferred theta phase on position and get a better estimate of its modulation by theta oscillations, we subtracted the phase precession fit from all spike theta phases in the place field. This returned deviation of theta phases from the best precession fit which was used to calculate precession adjusted TMI (TMIa) for each place field (Figure 4A). Since precession adjustment of TMI removed the influence of spatial location on the theta modulation index, it provided a better estimate of modulation of the neuron’s activity by theta oscillations. TMIa was higher than TMI for majority of place fields across dorsal CA1 (Figure 4B; 90 out of 102 place fields in pCA1 and 99 out of 115 place fields in dCA1, both proportions significantly greater than 50%; test for proportions; z = 7.72 and 7.74 for pCA1 and dCA1 respectively, p < 0.001 for both). Theta modulation measured using TMIa was statistically indistinguishable between pCA1 and dCA1 (Figure 4C; Wilcoxon rank sum test; z = 1.87, rank sum = 10730, p = 0.062; pCA1 median 0.848, dCA1 median 0.825, median difference 0.023 [−0.02, 0.06]; test for normality failed for post-hoc parametric tests, Lilliefors test p < 0.001 both for pCA1 and dCA1) with similar proportion of cells being significantly theta modulated after the adjustment for precession in the two sub-regions (Figure 4D; chi-square test for independence; χ^2^ = 0.01, p = 0.917, proportions of cells with significant TMIa in pCA1 = 91/97, dCA1 = 100/107 cells). Notice how the significant median difference in TMI (0.067 [0.002,0.143]; Figure 3) reported earlier in the results became non-significant after adjusting for phase precession (TMIa: median difference 0.023 [−0.02, 0.06]), indicating that there is no difference in the influence of theta oscillations on neural activity in pCA1 and dCA1.

**Figure 4.**
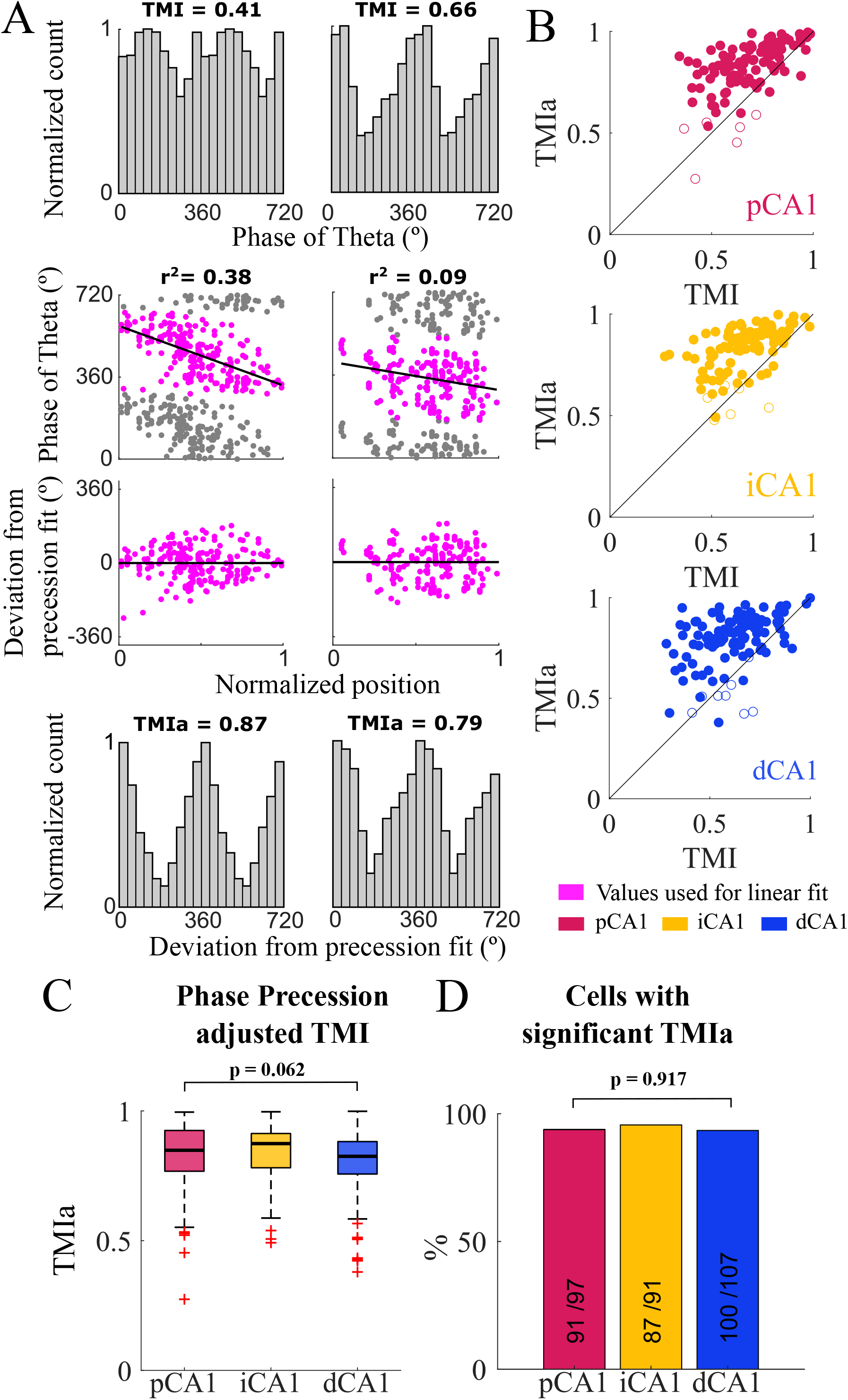
Phase precession adjusted TMI. **(A)** Examples showing estimation of theta modulation before and after adjusting for theta phase precession. Left: example place field with strong precession (statistically significant high r^2^) but low TMI. From top to bottom: spike phase histograms for all spikes in a place field, linear fit for phase precession, deviation of phases from the phase precession linear fit, and the histogram of these deviation angles used to calculate the phase precession adjusted TMI (TMIa). Right: place field of another cell with TMI close to the median TMI and weak precession (statistically significant low r^2^). **(B)** Scatter plots of TMIa vs. TMI. Each dot represents an individual place field; filled dots represent place fields with significant TMIa. Notice how most of the dots lie above the identity line in all sub-regions of CA1. **(C)** Box and whisker plots showing the distribution of TMIa for all place cells. **(D)** Proportion of cells with significant TMIa.

### Theta phase precession and modulation during MIS sessions

MIS sessions showed trends similar to the STD session. The strength of phase precession was statistically indistinguishable between pCA1 and dCA1 for all MIS sessions (Figure 5A; Wilcoxon rank sum test; p > 0.05; Supplementary Table 1). TMIa were higher than TMI for most place fields recorded in different MIS sessions (Figure 5B; proportions significantly greater than 50% for all MIS sessions; test for proportions, p < 0.001 for all MIS sessions for both pCA1 and dCA1; see Supplementary Table 1 for detailed statistics). TMIa also showed no significant difference between the two sub-regions in all MIS sessions (Figure 5C; Wilcoxon rank sum test; p > 0.05 for all comparisons; Supplementary Table 1). Similar proportions of place cells in pCA1 and dCA1 were significantly theta modulated during all MIS sessions (Figure 5D; chi-square test for independence; p > 0.05 for all MIS sessions except MIS 180° which showed p = 0.039; Supplementary Table 1).

**Figure 5.**
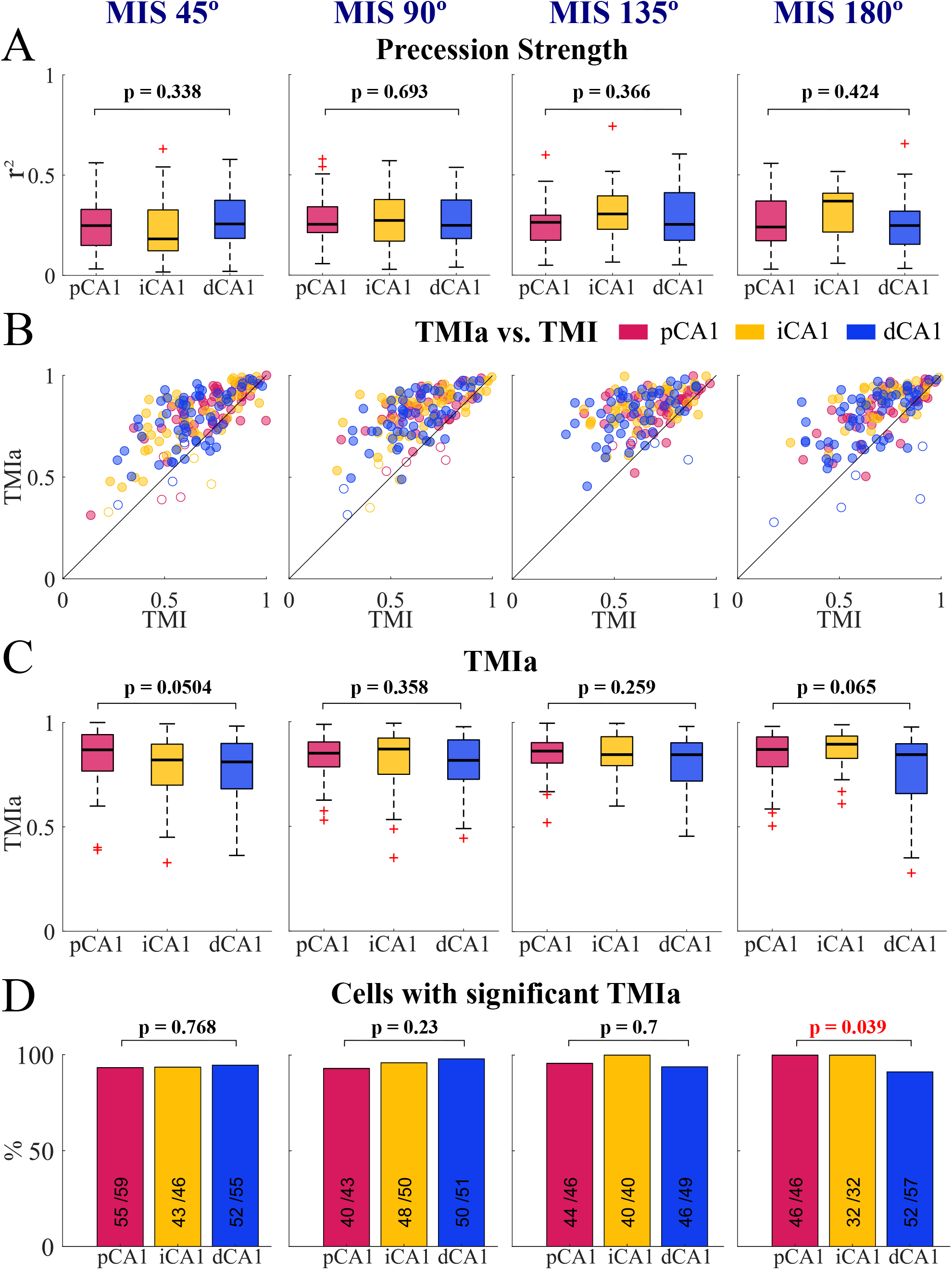
Theta phase coding during MIS sessions. Each column corresponds to the MIS angle shown at the top of each column. **(A)** Distribution of theta phase precession strength. **(B)** Scatter plots of TMIa vs. TMI. **(C)** Distribution of TMIa. **(D)** Proportion of cells with significant TMIa.

### Methodological concerns and statistical inference

Given that the negative results presented here differed from the previous results (Henriksen et al., 2010; Oliva et al., 2016), it is natural for the reader to wonder if methodological issues caused the loss of significance in this study. We performed several control analyses to ensure that that was not the case.

Difference in cluster isolation quality in pCA1 and dCA1 could affect the observed results. Hence, we explicitly tested for this possibility. There was no difference in the cluster isolation quality of the place cells from pCA1 and dCA1 included in all the analyses in this paper (Wilcoxon rank sum test, z = 0.66, rank sum = 10001, p = 0.51; 97 cells in pCA1 and 107 cells in dCA1).

The number of cells included in our study (97 cells in pCA1 + 91 cells in iCA1 + 107 cells in dCA1 = 295 cells) for the statistical comparison of phase precession and theta modulation were similar to the numbers reported by Oliva et al. (2016) (total 239 cells in pCA1, iCA1 and dCA1 together). Our numbers were much higher than the number of cells reported by Henriksen et al. (2010) (n = 38) for comparing theta modulation. Pooled standard deviation was 0.12 for strength of phase precession in our dataset. For this standard deviation, a sample size of n ≥ 34 in each group would be required to get a large effect size (Cohen’s d ≥ 0.8; Cohen, 1988, p25) and a statistical power of 90% in a two-sample t-test. This sample size requirement was satisfied in our comparisons. Thus, our tests have the statistical power to support the conclusion that pCA1 and dCA1 do not always exhibit functional differences.

To confirm the functional similarity along the CA1 transverse axis shown above, we also took a Bayesian approach to collect evidence for the alternate hypothesis that theta phase coding is different in the two sub-regions (H_1_: pCA1 ≠ dCA1), over the null hypothesis that it is not different between the two sub-regions in our paradigm (H_0_: pCA1 = dCA1). We calculated Bayes Factors (BF) to estimate if the data were more likely under H_0_ than H_1_. BF_10_ was < 1 for the strength of phase precession providing moderate evidence favoring H_0_ (BF_10_ = 0.158 for strength of precession). BF_10_ for TMIa was close to 1 (i.e. 1.07), suggesting both H_0_ and H_1_ were equally likely and so neither of the hypotheses could be supported. Thus, combining frequentist statistics and the Bayesian approach, we saw no theta-mediated functional differences along the CA1 transverse axis in our experimental protocol.

The cell populations of pCA1 and dCA1 included in all the statistical comparisons were not independently sampled in STD and different MIS sessions. As correction for multiple comparisons cannot be performed on samples that are not independent, the p values from single sessions here cannot be interpreted in isolation. However, trends across sessions are informative. The overall trend across STD and MIS sessions shows theta phase coding dynamics to be similar along the CA1 transverse axis in our experimental protocol. This observation is consistent with the previously experimental protocol (Deshmukh, 2021).

Some cells may have been repeatedly sampled across recording days (see methods for details). As a stringent control for repeat sampling, we only included cells from a day with maximum number of pyramidal cells recorded on a given tetrode. This reduced the number of place cells recorded in each CA1 sub-region across rats to about half (41 cells in pCA1 and 59 cells in dCA1, n = 11 rats) but satisfied the minimum requirement of n ≥ 34 in each group to see a large effect size with 90% power as calculated earlier in the results. The trends remained the same after controlling for repeat sampling. Theta modulation and theta phase precession were statistically indistinguishable between the two sub-regions (strength of phase precession: Wilcoxon rank sum test, p = 0.716; pCA1 median 0.289, dCA1 median 0.314, median difference −0.02 [−0.08,0.03] and TMIa: Wilcoxon rank sum test, p = 0.207; pCA1 median 0.86, dCA1 median 0.84, median difference 0.02 [−0.02,0.06]). The proportion of cells significantly theta modulated and with significant phase precession were also similar (chi-square test for independence, p = 0.48 and p = 0.32 for TMIa and strength of precession, respectively; 39/41 cells in pCA1 and 54/59 cells in dCA1 with significant TMIa; 40/41 cells in pCA1 and 55/59 cells in dCA1 with significant precession).

Theta oscillations are known to change in both amplitude and phase as a function of depth (Buzsáki, 2002). Theta amplitude is the lowest in the pyramidal cell layer and is the highest in the stratum lacunosum moleculare of CA1 (Buzsáki, 2002). Oliva et al. (2016) used a local reference electrode in the pyramidal cell layer of CA1 to estimate theta for showing significant difference in theta phase precession along the CA1 transverse axis. Despite this, there is a possibility that lower theta amplitude in the pyramidal cell layer affects theta phase estimation in our analyses. To eliminate this possibility, we identified tetrodes with recording locations below the cell layer (tetrode tracks terminating below the cell layer and above the hippocampal fissure; Supplementary Figure S3 A) in 7 out of 11 rats to be used as reference for theta phase estimation. All these tetrodes had higher theta power than that in the pyramidal cell layer (Supplementary Figure S3 B). We did not find significant differences in phase precession and TMIa between pCA1 and dCA1 in these rats (Supplementary Table 2) consistent with our results with local pyramidal cell layer theta as reference (Figure 2,3,4,5). The proportion of cells exhibiting theta phase coding were also statistically indistinguishable between pCA1 and dCA1 (Supplementary Table 2).

Next, we tested if the strength of local theta and proximodistal location of reference affected our results. For this, one pCA1 tetrode and one dCA1 tetrode with strongest theta in a recording session were chosen as references. Consistent with the results obtained from local theta reference (Figure 2,3,4,5), we found no significant difference in the strength of phase precession and TMIa between pCA1 and dCA1 using a pCA1 reference (Wilcoxon rank sum test, p > 0.05 for strength of phase precession and TMIa in STD and all MIS sessions) or a dCA1 reference (Wilcoxon rank sum test, p > 0.05 for strength of phase precession in STD and all MIS sessions; p > 0.05 for TMIa in all MIS sessions; borderline significance of TMIa, p = 0.041 only in STD session). Thus, the choice of the reference tetrode for theta phase estimation did not change the trend of theta modulation and theta phase precession along the CA1 transverse axis. pCA1 and dCA1 did not show theta-mediated functional differences in our experimental paradigm.

### LFP theta power is higher in dCA1 than pCA1

Slow oscillations in the LFP permit the integration of neural responses over long time windows and allow synchronization of neural activity over distributed brain regions (Colgin, 2013). The power of theta oscillations in LFP can be used to study the contribution of synchronized neural responses to the local network activity (Buzsáki, 2002; Buzsáki et al., 2012; Colgin, 2013). To check if pCA1 and dCA1 networks showed differences in the strength of theta synchronized local activity, we looked at total power in the theta frequency range along the CA1 transverse axis. Theta power and theta phase are known to change along with the depth of the electrode within CA1 (Buzsáki, 2002). Tetrode recordings do not allow simultaneous measurement of precise layer-wise LFP power unlike linear probes. However, comparing theta power at matching depths across the proximodistal axis mitigates the influence of tetrode depth on theta power. To ensure that the LFPs included were from the same depth of CA1, we analyzed LFP theta power and theta peak frequency only for the tetrodes which were in the pyramidal cell layer on the day of recording, as confirmed from the putative pyramidal cell spiking activity. The power spectrum of each tetrode, after removing the 1/ f^n^ component, was normalized by the highest power in the theta range across all tetrodes within a rat. Theta peak frequency and total power in the theta range for each region (pCA1, iCA1 and dCA1) for the given rat was determined from the means of the normalized power spectra of all tetrodes in the given region (Figure 6A, S4). To avoid oversampling the same region, data from a day with the maximum number of tetrodes recording pyramidal cell activity during a session was chosen from each rat to perform the statistics.

**Figure 6.**
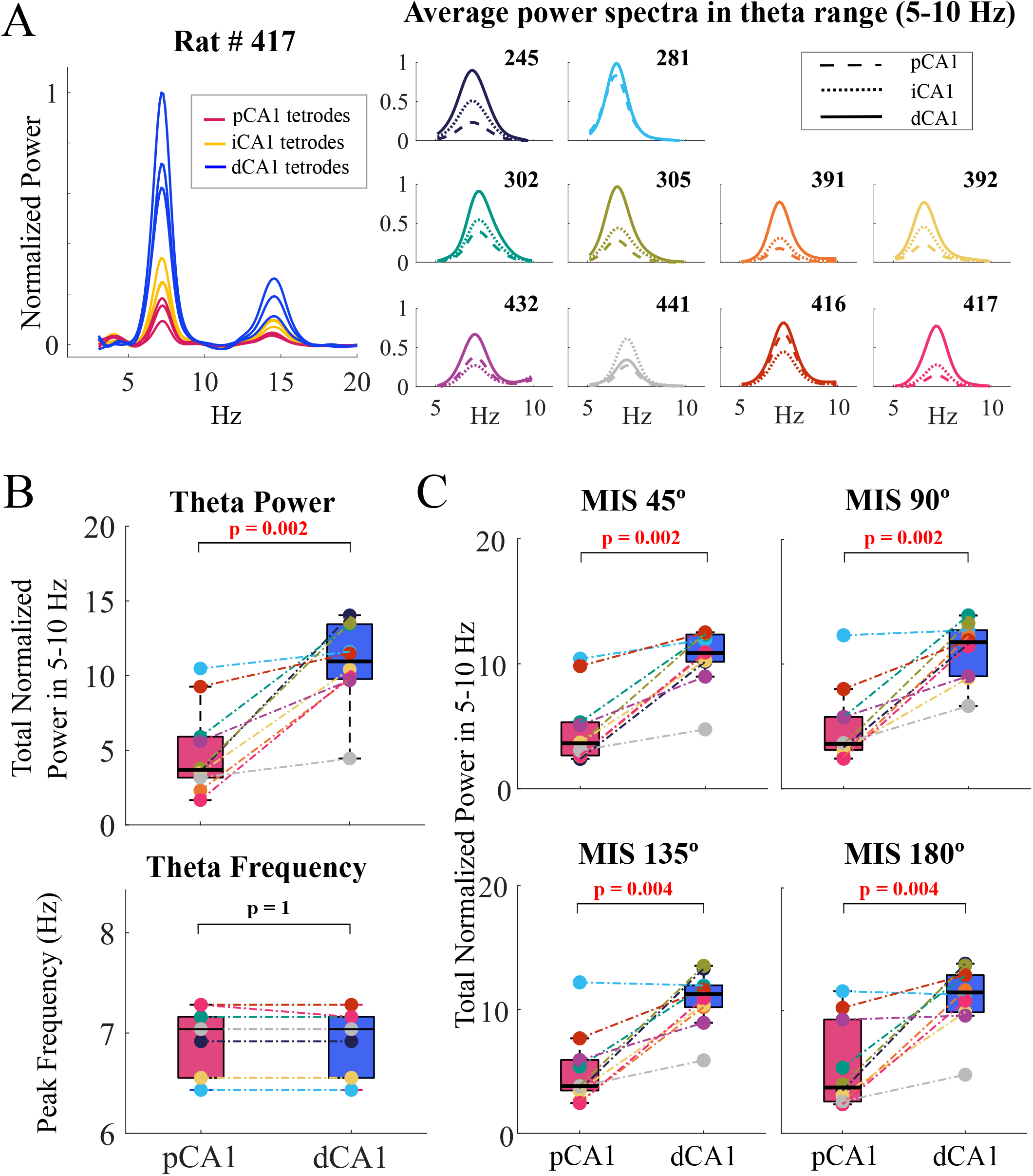
Theta power along the CA1 transverse axis. **(A)** Left: example power spectra of individual tetrodes from one rat normalized by the maximum power in the theta range. Right: average power spectra in the theta range of pCA1, iCA1, and dCA1 tetrodes for each rat for STD session **(B)** Total power in the theta range (top) and peak theta frequency (bottom) in STD session across all rats (rats represented as dots by the same color as in the right plot in A) **(C)** Total power in the theta range for MIS sessions. Across STD and MIS sessions, dCA1 LFPs tended to have higher power in theta range than pCA1 LFPs.

Out of the 11 rats included in this study, 10 had tetrodes simultaneously recording pyramidal cell activity in both pCA1 and dCA1. In these 10 rats, we found that dCA1 had higher theta power than pCA1 in the STD session (Figure 6B; Wilcoxon signed rank test; W = 0, p = 0.002; pCA1 median 3.7, dCA1 median 11, median difference −7.3 [−10.2, −4.7]; test for normality passed for post-hoc parametric tests, Lilliefors test p = 0.07 and 0.1 for pCA1 and dCA1, respectively; Post-hoc paired t-test, t (9, 3.4) = −5.4, p < 0.001; Cohen’s d = −1.9 [−3, −0.9]; n = 10 rats). The proportion, 10/10 rats, was significantly greater than the chance proportion of 0.5 (test for proportions, z = 3.16, p < 0.001). The peak theta frequencies in pCA1 and dCA1 were statistically indistinguishable (Figure 6B; Wilcoxon signed rank test; W = 1, p = 1; pCA1 median 7.04 Hz, dCA1 median 7.04 Hz, median difference 0 [−0.5,0.5] Hz). A similar trend of higher theta power in dCA1 than pCA1 was observed in all MIS sessions (Figure 6C; Wilcoxon signed rank test; p < 0.01 for all MIS sessions, see Supplementary Table 1). 10/11 rats were included for all MIS sessions, after ensuring every rat had at least one tetrode each in the pCA1 and dCA1 pyramidal cell layer. All rats continued to show greater theta power in dCA1 than pCA1 in MIS 45° and 90° sessions (the proportion 10/10 rats is significantly greater than the chance proportion of 0.5; test for proportions, z = 3.16, p < 0.001); all except one rat continued to show greater theta power in dCA1 than pCA1 in MIS 135°and 180° sessions (the proportion 9/10 rats is significantly greater than the chance proportion of 0.5; test for proportions, z = 2.53, p = 0.006). The peak theta frequencies in pCA1 and dCA1 were statistically indistinguishable in all MIS sessions (statistics in Supplementary Table 1).

Theta oscillation amplitude and frequency in different parts of the hippocampus and entorhinal cortex are known to vary with the animal’s speed (Bland & Oddie, 2001; Hinman et al., 2011; Long et al., 2014). Speed modulation of CA1 theta amplitude and theta frequency was observed in our experimental protocol but did not show any significant differences between pCA1 and dCA1 (Supplementary Figure S5).

## Discussion

### Comparison with the previous reports of theta dynamics and spatial selectivity along the transverse axis of CA1

Based on anatomical connectivity patterns, MEC is thought to be a part of the dorsal “where” stream, while LEC is thought to be a part of the ventral “what” stream (Mishkin et al., 1983; Suzuki & Amaral, 1994; Knierim et al., 2014). MEC and LEC project differentially along the transverse axis of pCA1 (Steward & Scoville, 1976; Witter & Amaral, 2004; Ito & Schuman, 2011; Masurkar et al., 2017). Correlated with strong theta modulated path integration based inputs from MEC, higher spatial selectivity and theta modulation have been shown in pCA1 compared to dCA1 (Henriksen et al., 2010; Igarashi et al., 2014; Oliva et al., 2016).

In this paper, we have demonstrated that pyramidal cells in dCA1 and pCA1 show comparable theta phase precession and theta modulation when the rats run on a circular track with 4 distinct local textures in the presence of multiple global cues. An identical proportion of spatially selective cells in dCA1 and pCA1 exhibit theta phase precession and theta modulation suggesting comparable levels of theta mediated activation along the transverse axis of CA1 in representing this cue-rich environment. In addition, dCA1 shows higher power in LFP theta oscillations than pCA1. This work complements our earlier report that in the same experimental protocol, dCA1 and pCA1 have comparable spatial selectivity (Deshmukh, 2021). These results indicate that the previously reported lower spatial selectivity and correspondingly lower theta modulation and theta phase precession in dCA1 than pCA1 (Henriksen et al., 2010; Oliva et al., 2016) are not immutable, and experimental conditions and/or task demands may alter the balance.

Spatial selectivity in LEC and dCA1 has previously been shown to be influenced by experimental conditions. LEC shows spatial selectivity in the presence of objects (Deshmukh & Knierim, 2011) and shows representation of past locations of objects (Deshmukh & Knierim, 2011; Tsao et al., 2013), but does not show spatial selectivity in the absence of objects (Hargreaves et al., 2005). Consistent with this, dCA1 has been shown to be more spatially selective in the presence of objects (Burke et al., 2011; Vandrey et al., 2021). However, the increased spatial selectivity in dCA1 may not always be inherited from LEC or dependent on presence of objects. In the present experimental protocol, LEC is non-selective to space (Yoganarasimha et al., 2011) and has very weak theta power and theta modulation (Deshmukh et al., 2010). Thus, the change in the functionality of dCA1 from the previously reported simple environments to the more complex environment reported here cannot be explained as being caused by stronger sensory derived spatial and theta modulated/precessing inputs from LEC in the presence of distinct textures on the track.

Responses of pCA1 and dCA1 to the cue-mismatch manipulations need to be considered before proposing plausible mechanisms for the above results. Theta modulation and theta phase precession were found to be comparable between pCA1 and dCA1 for almost all cue configurations across rotates coherently with global cues from STD to MIS sessions, while pCA1 representation loses coherence (Deshmukh, 2021). Loss of coherence in pCA1 can be explained by strong competing inputs it receives – MEC representation rotates with global cues (Neunuebel et al., 2013), while distal CA3 representation rotates with local cues (Lee et al., 2015). In contrast, neither LEC (Neunuebel et al., 2013) nor proximal CA3 (Lee et al., 2015) show coherent rotation with global cues. We (Deshmukh, 2021) proposed that in the absence of strong competing inputs, the less numerous inputs from MEC (Neunuebel et al., 2013; Masurkar et al., 2017), subiculum (Ding, 2013; Xu et al., 2016; Sharma et al., 2022), and nucleus reuniens (Dolleman-Van Der Weel & Witter, 1996; Vertes et al., 2006; Jankowski et al., 2014, 2015; Dolleman-van der Weel et al., 2019) may be sufficient to drive dCA1 to rotate coherently with the global cues. While these inputs might be sufficient to explain the responses along the transverse axis of CA1 to cue mismatch, additional mechanisms must be considered to explain comparable spatial selectivity in pCA1 and dCA1 in our task (Deshmukh, 2021) in contrast to the previous reports (Henriksen et al., 2010; Oliva et al., 2016). Spatial selectivity of the MEC/subiculum/nucleus reuniens inputs may be higher in our complex environment than in the previous simple environments. Alternatively, excitability in dCA1 may increase in response to higher task demand (Deshmukh, 2021).

Can these mechanisms proposed to explain the patterns of spatial selectivity and coherence of responses to cue manipulation along the transverse axis of CA1 also explain the difference in theta phase precession and modulation between the previous reports (Henriksen et al., 2010; Oliva et al., 2016) and the present study? Spatially selective inputs from MEC/subiculum/nucleus reuniens are theta modulated (Cacucci et al., 2004; Hafting et al., 2008; Tsanov et al., 2011; Jankowski et al., 2014, 2015; Poulter et al., 2021). An increase in their theta modulation and/or spatially selective response amplitude may result in higher theta responsivity in dCA1 in the present task. Alternatively, an increase in theta drive onto dCA1 from the pacemaker, the medial septum (Petsche et al., 1962; Gaykema et al., 1990; Bland & Oddie, 2001; Koenig et al., 2011), in the present task may explain our results. To the best of our knowledge, the medial septum does not show differential projection patterns along the transverse axis of CA1, and we do not know if the pacemaker inputs to pCA1 and dCA1 can be differentially modulated. Thirdly, neuromodulatory inputs may increase the excitability of dCA1 neurons in the present task compared to the previous reports (Henriksen et al., 2010; Oliva et al., 2016). Such increase in excitability can lead to both increased spatial selectivity as well as increased theta oscillatory dynamics in dCA1 by making it more responsive to inputs from MEC/subiculum/nucleus reuniens and the medial septum, eliminating the need for the differential 2011) are not theta modulated (Deshmukh et al., 2010) and rotate weakly with the local cues (Neunuebel et al., 2013) in the present task. These LEC inputs cannot explain the coherent rotation of dCA1 with global cues. However, higher firing rates of LEC neurons in the present task compared to the previous reports may lead to increased theta oscillatory dynamics as well as spatial selectivity in dCA1. Higher activity of nonspatial non-theta-modulated LEC inputs may lead to the dCA1 neurons being closer to the action potential threshold, effectively increasing their excitability, and thus giving rise to more robust responses to spatially selective as well as theta oscillatory inputs. Experiments explicitly comparing the activity of pCA1 and dCA1 neurons as well as their inputs in environments of different levels of complexity and task demands, beyond the scope of the present work, are necessary to disambiguate between these plausible mechanisms.

### LFP theta power in pCA1 and dCA1

LFPs in dCA1 show stronger theta oscillations than pCA1 in our experimental protocol. Previous studies (Henriksen et al., 2010; Oliva et al., 2016) did not compare amplitudes of theta oscillations in pCA1 and dCA1 LFPs. Hence, we do not know if the relative theta power in dCA1 and pCA1 changes from the previous studies to the present study. The observed difference in the amplitude of the LFP theta oscillations between pCA1 and dCA1 could be caused by a variety of factors such as morphology and density of the pyramidal cells, laminar organization and density of their inputs, local curvature, etc. (Buzsáki, 2002; Buzsáki et al., 2012). Speed modulation of theta oscillations is comparable in pCA1 and dCA1. Thus, the difference in theta power in LFP may not be caused by pCA1 and dCA1 theta generators being differentially modulated by speed. Higher theta power in dCA1 LFPs than pCA1 LFPs, together with comparable strength of theta modulation between dCA1 and pCA1 neurons, suggests that theta oscillations can drive the dCA1 network at least as strongly as the pCA1 network under certain task conditions.

### Summary

This study, together with Deshmukh (2021), challenges the notion that dCA1 is inherently less spatial and has weaker theta oscillatory dynamics than pCA1. We suggest that LEC and MEC drives onto dCA1 and pCA1 respectively are not nearly as mutually exclusive as suggested earlier, and their relative contributions may change based on task demands. Further experiments are necessary to transverse axis of CA1 and their task dependent modulation.

## Acknowledgements

We thank Jeremy Johnson, Geeta Rao, Vyash Puliyadi, Amanda Smolinsky, Lou Blanpain, and Kimberley Christian for their support in data collection; Bruce L. McNaughton for discussion; Rajat Saxena and Indraja R. Jakhalekar for inputs on the manuscript. This work was supported by Wellcome Trust/DBT India Alliance Grant IA/S/13/2/501024 and Pratiksha trust grant PE/CHAIR-19-025.03. Data collection was supported by R01 NS039456 (J. Knierim, PI).

