## Supplementary Information for "Comparable theta phase coding dynamics along the transverse axis of CA1"

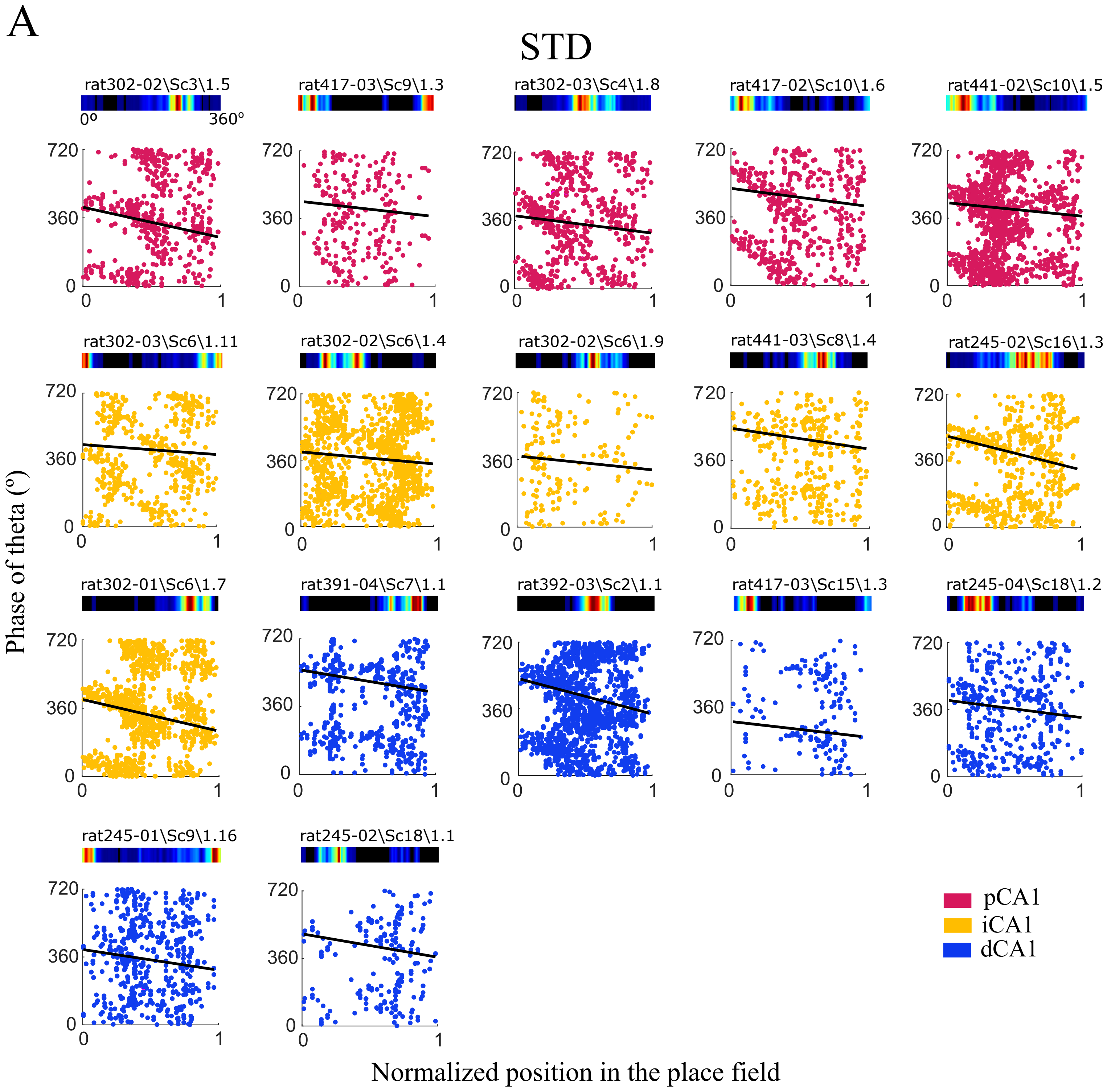
A
STD
rat302-02\Sc3\1.5
0º
360°
rat417-03\Sc9\1.3
rat302-03\Sc4\1.8
rat417-02\Sc10\1.6
rat441-02\Sc10\1.5
rat302-03\Sc6\1.11
rat302-02\Sc6\1.4
rat302-02\Sc6\1.9
rat441-03\Sc8\1.4
rat245-02\Sc16\1.3
rat302-01\Sc6\1.7
rat391-04\Sc7\1.1
rat392-03\Sc2\1.1
rat417-03\Sc15\1.3
rat245-04\Sc18\1.2
rat245-01\Sc9\1.16
rat245-02\Sc18\1.1
720
360
0
1
Phase of theta (º)
Normalized position in the place field
pCA1
iCA1
dCA1

Figure S1(A)

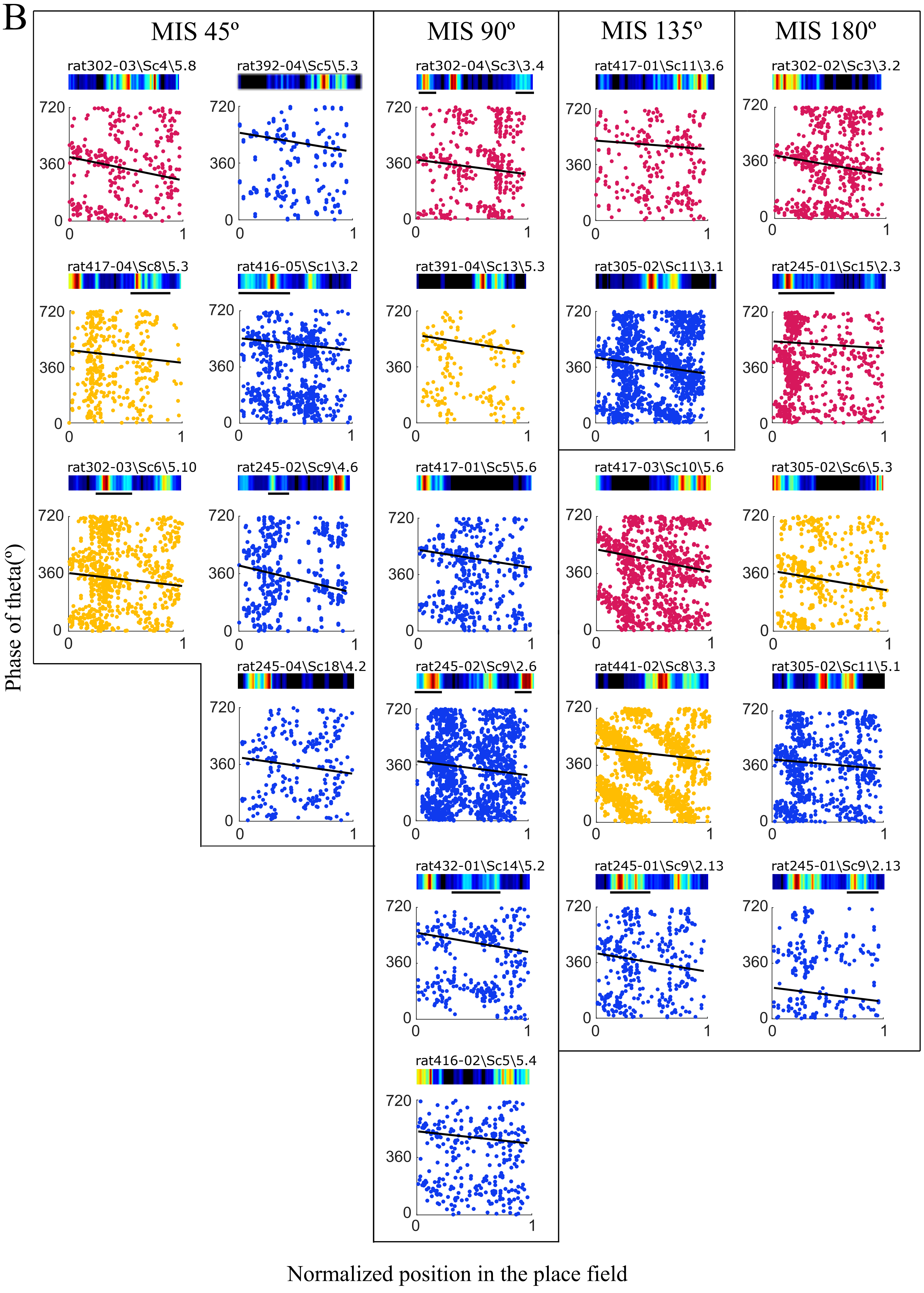

B
MIS 45º
MIS 90º
MIS 135º
MIS 180º
rat302-03\Sc4\5.8
rat392-04\Sc5\5.3
rat302-04\Sc3\3.4
rat417-01\Sc11\3.6
rat302-02\Sc3\3.2
rat417-04\Sc8\5.3
rat416-05\Sc1\3.2
rat391-04\Sc13\5.3
rat305-02\Sc11\3.1
rat245-01\Sc15\2.3
rat302-03\Sc6\5.10
rat245-02\Sc9\4.6
rat417-01\Sc5\5.6
rat417-03\Sc10\5.6
rat305-02\Sc6\5.3
rat245-04\Sc18\4.2
rat245-02\Sc9\2.6
rat441-02\Sc8\3.3
rat305-02\Sc11\5.1
rat432-01\Sc14\5.2
rat245-01\Sc9\2.13
rat245-01\Sc9\2.13
rat416-02\Sc5\5.4
720
360
0
1
Phase of theta(º)
Normalized position in the place field


Figure S1(B)

**Figure S1. Cells with multiple theta phase precession cycles per place field.** Few place fields showed more than one cycle of phase precession, indicating that these probably were multiple overlapping place fields (Skaggs et al., 1996). These place fields were removed manually from the analysis of theta phase precession and theta modulation to avoid confounds due to the overlapping precession cycles. This figure shows all place fields excluded for this reason. For each place field, top: linearized rate map of the place cell over the entire track (0°- 360°), black represents 0 Hz firing rate, blue to red depict low to high firing rates. Bottom: precession of spike theta phase with position of the animal in the place field. Zero marks the start, and 1 marks the end of the place field. Red: spike theta phases for place fields in pCA1, yellow: iCA1, blue: dCA1, black line: linear regression fit showing a bad fit caused by attempting to fit multiple precession cycles with a single line. For cells with more than one identified place field, only the field that was removed from the analysis has been shown here. These fields are marked with a line under the rate map. (**A**) Cells removed in STD session (**B**) Cells removed in MIS sessions.

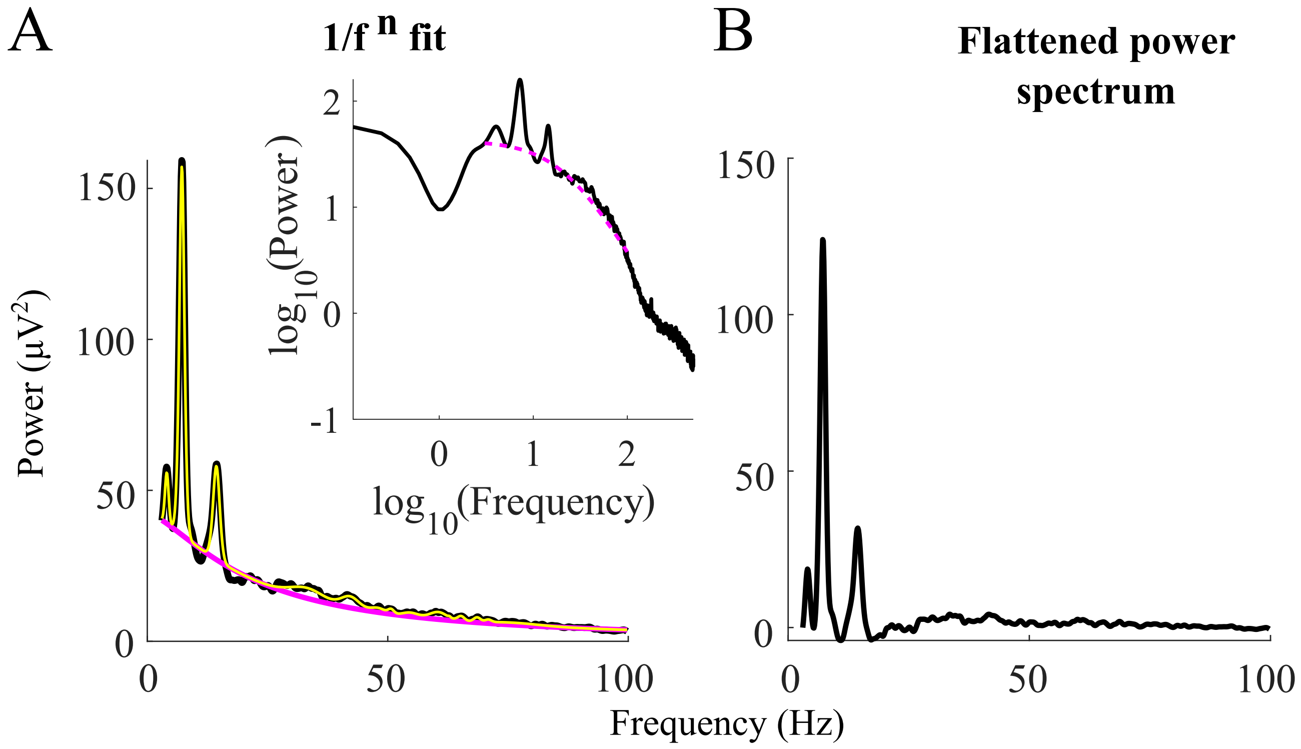


**Figure S2. Subtraction of 1/f$^n$ using FOOOF. (A)** Example power spectrum showing the 1/f$^n$ property of LFP. Inset shows the power spectrum plotted in the log-log scale showing that the 'knee' parameter of the FOOOF algorithm used over 3-100 Hz controls for the bend in the 1/f$^n$ component. Black trace: original power spectrum, pink dotted trace: 'background' 1/f$^n$ fit, yellow trace: power spectrum modelled by the algorithm after accounting for the 1/f$^n$ background and Gaussian peaks. **(B)** 'Flattened' spectrum obtained after subtracting the 'background' fit (pink) from the original spectrum (black).

Figure S2

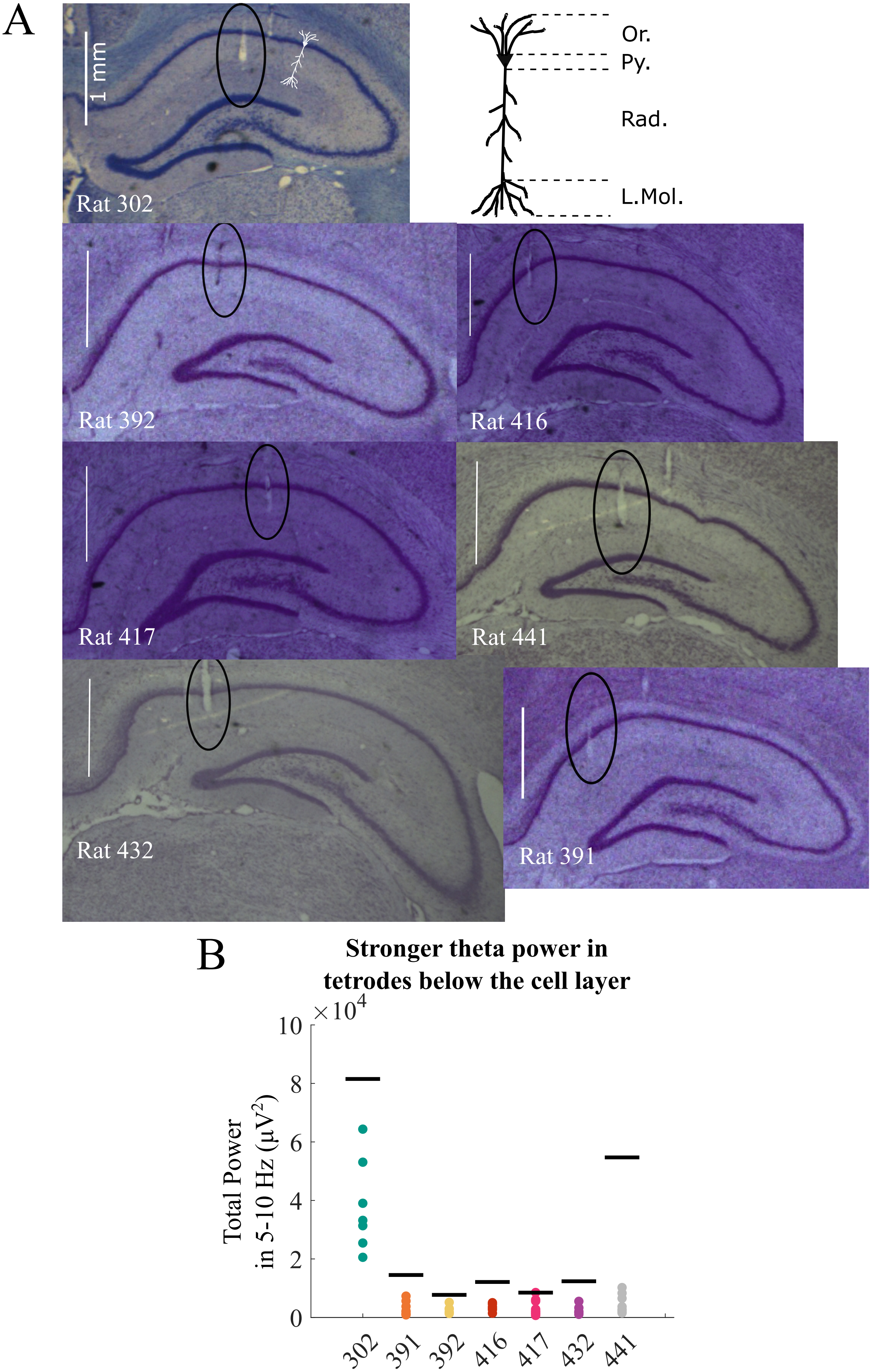
A
1 mm
Rat 302
Or.
Py.
Rad.
L.Mol.
Rat 392
Rat 416
Rat 417
Rat 441
Rat 432
Rat 391
B
Stronger theta power in
tetrodes below the cell layer
$\times 10^4$
10
8
6
4
2
0
Total Power
in 5-10 Hz ($\mu V^2$)
302
391
392
416
417
432
441

Figure S3

**Figure S3. Theta reference below the cell layer. (A)** Recording locations of tetrodes chosen as theta reference to control for depth-wise differences in theta oscillations influencing phase estimation. Tetrodes recording LFP below the cell layer (tetrode tracks marked with black circles) could be identified in 7 out of 11 rats. A single reference tetrode was used for all CA1 cells in a rat. Illustration of a neuron marks the different layers of CA1. Location below the cell layer was based on histology and the power of theta oscillations in the LFP **(B)** The chosen theta references had higher LFP theta power in each rat than all tetrodes in the pyramidal cell layer. Black lines represent theta power of the chosen reference as marked in A and positioned between Rad. and L.Mol., Colored dots represent tetrodes that recorded pyramidal cell activity and were confirmed to be in Py. Or: stratum oriens, Py: stratum pyramidale, Rad: stratum radiatum, L.Mol.: stratum lacunosum moleculare.

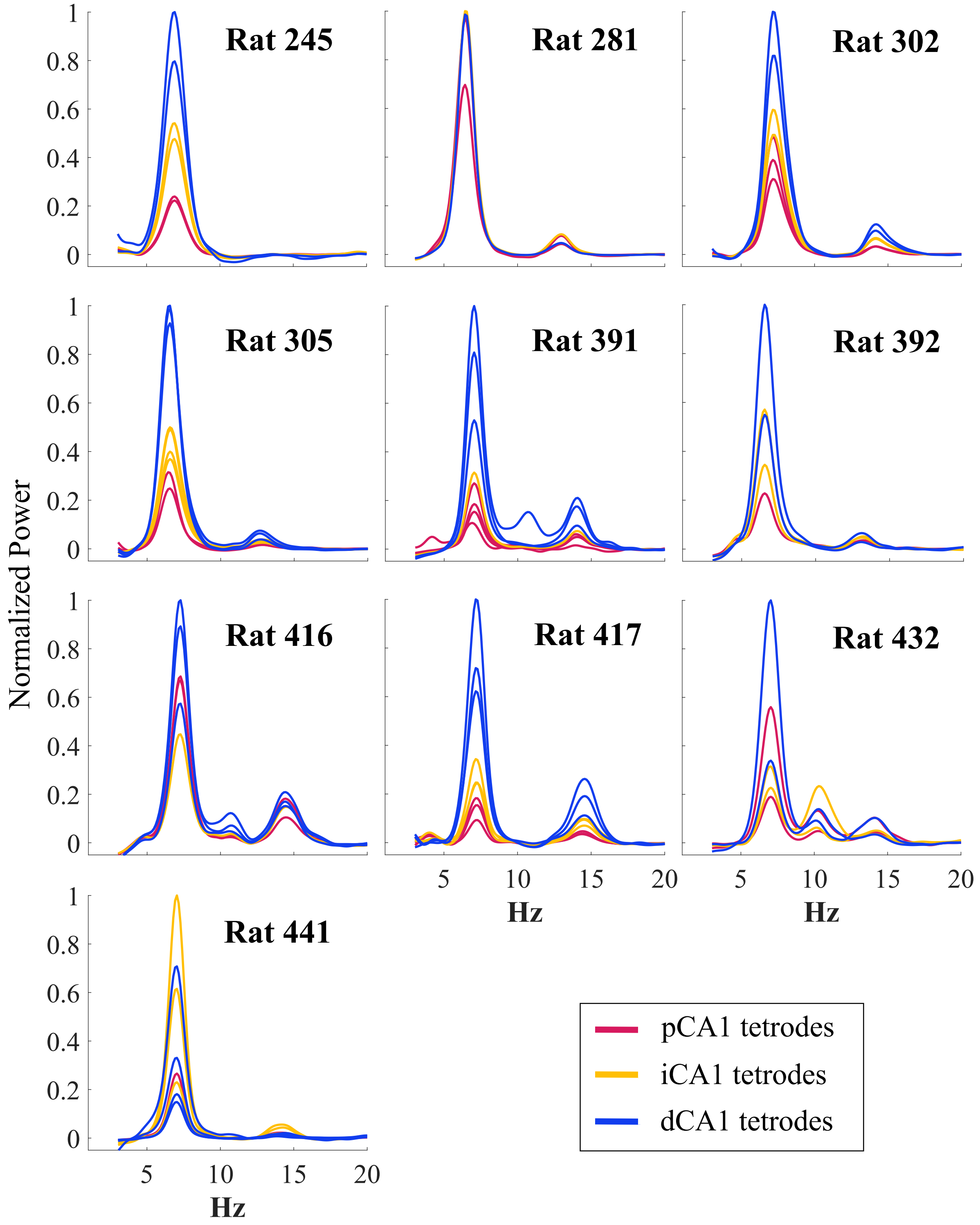


**Figure S4. LFP theta power for STD session.** Power spectra of individual tetrodes along the CA1 transverse axis for all animals used in the study. The $1/f^n$ component of the spectrum was removed for each tetrode using the FOOOF algorithm (Figure S2).

Figure S4

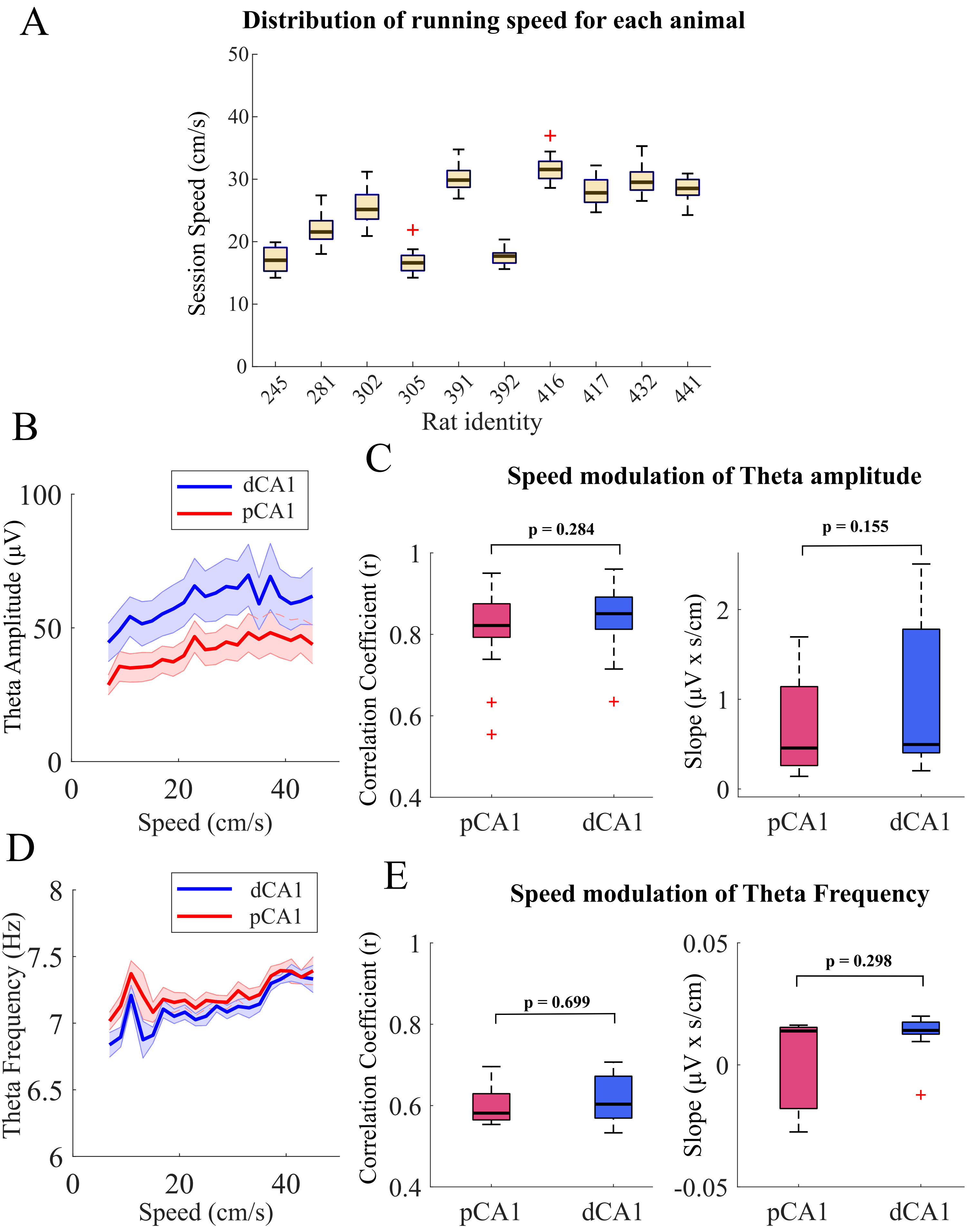
A
Distribution of running speed for each animal
Session Speed (cm/s)
50
40
30
20
10
0
245
281
302
305
391
392
416
417
432
441
Rat identity
B
dCA1
pCA1
Theta Amplitude (μV)
100
50
0
0
20
40
Speed (cm/s)
C
Speed modulation of Theta amplitude
p = 0.284
Correlation Coefficient (r)
1
0.8
0.6
0.4
pCA1
dCA1
p = 0.155
Slope (μV x s/cm)
2
1
0
pCA1
dCA1
D
dCA1
pCA1
Theta Frequency (Hz)
8
7.5
7
6.5
6
0
20
40
Speed (cm/s)
E
Speed modulation of Theta Frequency
p = 0.699
Correlation Coefficient (r)
1
0.8
0.6
0.4
pCA1
dCA1
p = 0.298
Slope (μV x s/cm)
0.05
0
-0.05
pCA1
dCA1

Figure S5

**Figure S5. Speed modulation of LFP theta oscillation.** Modulation of LFP theta amplitude by the rat's speed was analyzed to test if the difference in the LFP theta power between pCA1 and dCA1 (Figure 6) was correlated with differential modulation of LFP theta oscillation by the animal's running speed. Speed was estimated between each consequent frame (~33 ms) and Gaussian smoothed (window size = 5 sample points, i.e., ~165 ms, $\sigma = 1$). To remove points where the animal was sitting and points of erroneous speed calculation due to occlusion of some of the tracking LEDs, timestamps with speeds ≤ 2 cm/s and ≥ 150 cm/s were discarded. Mean speed was calculated in every 1 s non-overlapping window. Theta amplitude in each window was calculated as the mean envelope of the signal filtered in the theta range (5-10 Hz) and theta frequency was calculated as the mean time difference between the consecutive peaks of theta cycles detected in the window. **(A)** Box and whisker plots showing the distribution of session speeds for each animal. Session speed was calculated as the mean running speed of the animal in a recording session. Session speeds ranged from 10 cm/s to 40 cm/s across all animals. **(B)** Theta amplitude increased with increasing running speed in STD session for both dCA1 and pCA1 (n = 10 rats, shaded regions represent standard error of the mean). **(C)** Distributions of correlation coefficient (r) between theta amplitude and speed (**left**) and slope of linear fit (**right**) were comparable between pCA1 and dCA1 (Wilcoxon rank sum test, z = -1.07, rank sum = 387, p = 0.28 for correlation coefficient; z = -1.42, rank sum = 373, p = 0.15 for slope). Linear regression fits were performed for theta amplitude vs. animal speed and theta frequency vs. animal speed only in the 10-40 cm/s range after binning the speeds in 2 cm/s wide bins, as there were substantially fewer samples available at speeds higher than 40 cm/s and lower than 10 cm/s, compared to the samples in the 10-40 cm/s range. Only the tetrodes with significant linear fits ($p < 0.05$) were included in the statistics for correlation coefficient (r) and slope comparisons between pCA1 and dCA1. Around 90% of the tetrodes in both pCA1 and dCA1 across 10 rats for STD session showed significant linear fits between theta power and speed (20/22 tetrodes in pCA1 and 22/25 tetrodes in dCA1). **(D), (E)** Same as (B) and (C) but for theta frequency. Theta frequency did not show significant differences in speed modulation between pCA1 and dCA1 (Wilcoxon rank sum test for $n < 10$; rank sum = 34, p = 0.69 for correlation coefficient, rank sum = 29, p = 0.29 for slope). Only a small fraction of the tetrodes in both pCA1 and dCA1 across 10 rats for STD session showed significant linear fits between theta frequency and speed (5/22 tetrodes in pCA1 and 9/25 tetrodes in dCA1).

**Supplementary table 1:** Statistical comparisons for MIS sessions

| **Comparison of strength of phase precession between pCA1 and dCA1** | | | | | |
|---|---|---|---|---|---|
| | | **MIS 45°** | **MIS 90°** | **MIS 135°** | **MIS 180°** |
| pCA1 median | | 0.248 | 0.253 | 0.26 | 0.24 |
| dCA1 median | | 0.256 | 0.248 | 0.25 | 0.247 |
| Median difference [95% CI] | | -0.008 [-0.08, 0.05] | 0.005 [-0.04, 0.05] | 0.01 [-0.07, 0.07] | -0.007 [-0.09, 0.09] |
| Wilcoxon rank sum test | z value | -0.958 | 0.394 | -0.905 | 0.8 |
| | rank sum | 3223 | 2095 | 2086 | 2513 |
| | p value | 0.338 | 0.693 | 0.366 | 0.424 |
| Post-hoc Lilliefor's test | p value (pCA1) | 0.85 | 0.003 | 0.097 | 0.018 |
| | p value (dCA1) | 0.41 | 0.053 | 0.23 | 0.078 |
| Post-hoc t-test | (df*, σ) | (112, 0.13) | Failed test for normality | (93, 0.13) | Failed test for normality |
| | p value | 0.269 | | 0.263 | |
| **Comparison of proportion of cells with significant precession between pCA1 and dCA1 (chi-square test for independence)** | | | | | |
| $\chi^2$ value | | 0.038 | 0.015 | 0.15 | 0.605 |
| p value | | 0.845 | 0.903 | 0.698 | 0.436 |
| proportion of cells in pCA1 | | 53/59 | 42/43 | 44/46 | 44/46 |
| proportion of cells in dCA1 | | 50/55 | 50/51 | 46/49 | 56/57 |
| **Place fields with TMIa greater than TMI (test for proportions)** | | | | | |
| **pCA1** | | | | | |
| z value | | 5.5 | 4.42 | 5.65 | 4.3 |
| p value | | < 0.001 | < 0.001 | < 0.001 | < 0.001 |
| proportion of place fields in pCA1 | | 54/64 | 38/46 | 45/50 | 40/49 |
| **dCA1** | | | | | |
| z value | | 5.16 | 6.49 | 6.38 | 1.6 |
| p value | | < 0.001 | < 0.001 | < 0.001 | < 0.001 |
| proportion of place fields in dCA1 | | 50/60 | 53/57 | 49/52 | 52/63 |
| **Comparison of TMIa between pCA1 and dCA1 (Wilcoxon rank sum test)** | | | | | |
| pCA1 median | | 0.87 | 0.84 | 0.863 | 0.87 |
| dCA1 median | | 0.81 | 0.81 | 0.846 | 0.85 |
| Median difference [95% CI] | | 0.06 [-0.02, 0.12] | 0.03 [-0.02, 0.1] | 0.017 [-0.04, 0.07] | 0.02 [-0.04, 0.11] |
| Wilcoxon rank sum test | z value | 1.956 | 0.918 | 1.128 | 1.847 |
| | rank sum | 3738 | 2164 | 2360 | 2671 |
| | p value | 0.0504 | 0.358 | 0.259 | 0.065 |

| | | | | | |
|---|---|---|---|---|---|
| Post -hoc Lilliefor's test | p value (pCA1) | 0.033 | 0.076 | 0.19 | 0.039 |
| | p value (dCA1) | 0.012 | 0.259 | 0.055 | < 0.001 |
| Post-hoc t-test | (df, σ) | Failed test for normality | (92, 0.12) | (93, 0.1) | Failed test for normality |
| | p value | | 0.298 | 0.164 | |
| **Comparison of proportion of cells with significant TMIa between pCA1 and dCA1 (chi-square test for independence)** | | | | | |
| $\chi^2$ value | | 0.087 | 1.44 | 0.15 | 4.241 |
| p value | | 0.768 | 0.23 | 0.698 | 0.039 |
| proportion of cells in pCA1 | | 55/59 | 40/43 | 44/46 | 46/46 |
| proportion of cells in dCA1 | | 52/55 | 50/51 | 46/49 | 52/57 |
| **Comparison of total power in theta range between pCA1 and dCA1 (Wilcoxon signed rank test)** | | | | | |
| pCA1 median | | 3.642 | 3.61 | 3.813 | 3.73 |
| dCA1 median | | 10.85 | 11.75 | 11.23 | 11.47 |
| Median difference [95% CI] (normalized power) | | -7.2 [-9.3, -4.5] | -8.14 [-9.7, -5.4] | -7.4 [-9.7, -4.9] | -7.7 [-10.2, -2] |
| signed rank | | 0 | 0 | 1 | 1 |
| p value | | 0.002 | 0.002 | 0.004 | 0.004 |
| Post -hoc Lilliefor's test | p value (pCA1) | 0.07 | 0.03 | 0.052 | 0.055 |
| | p value (dCA1) | 0.09 | 0.07 | 0.41 | 0.43 |
| Post-hoc paired t-test | (df, σ) | (9, 2.88) | Failed test for normality | (9, 3.4) | (9, 4.06) |
| | p value | < 0.001 | | < 0.001 | 0.0017 |
| **Comparison of peak theta frequency between pCA1 and dCA1 (Wilcoxon signed rank test)** | | | | | |
| pCA1 median | | 7.09 | 6.92 | 6.97 | 6.85 |
| dCA1 median | | 7.09 | 6.92 | 6.91 | 6.85 |
| Median difference [95% CI] (Hz) | | 0 [-0.7, 0.6] | 0 [-0.5, 0.5] | 0.06 [-0.5, 0.5] | 0 [-0.3, 0.3] |
| signed rank | | 6 | 0 | 7 | 0 |
| p value | | 0.875 | 1 | 0.625 | 1 |
| Post -hoc Lilliefor's test | p value (pCA1) | 0.148 | 0.4 | 0.33 | 0.316 |
| | p value (dCA1) | 0.434 | 0.5 | 0.5 | 0.242 |
| Post-hoc paired t-test | (df, σ) | (9, 0.076) | (9, 0.038) | (9, 0.081) | (9, 0.038) |
| | p value | 0.34 | 0.34 | 1 | 0.34 |

(*df = degrees of freedom, σ = standard deviation)

**Supplementary table 2:** Statistics for pCA1 and dCA1 using single theta reference below the cell layer

| **Strength of phase precession for all place cells** | | | | | | |
|---|---|---|---|---|---|---|
| | | **STD1** | **MIS 45°** | **MIS 90°** | **MIS 135°** | **MIS 180°** |
| pCA1 median | | 0.302 | 0.267 | 0.264 | 0.28 | 0.252 |
| dCA1 median | | 0.338 | 0.265 | 0.28 | 0.32 | 0.238 |
| Median difference [95% CI] | | -0.03 [-0.09, 0.02] | 0.002 [-0.07, 0.06] | -0.016 [-0.09, 0.06] | -0.04 [-0.12, 0.08] | 0.0124 [-0.08, 0.12] |
| Wilcoxon rank sum test | z value | -1.252 | -1.07 | -0.799 | -1.22 | 0.418 |
| | rank sum | 5920 | 2233 | 1089 | 1184 | 1558 |
| | p value | 0.211 | 0.283 | 0.424 | 0.222 | 0.675 |
| Post-hoc Lilliefor's test | p value (pCA1) | 0.99 | 0.24 | 0.049 | 0.364 | 0.178 |
| | p value (dCA1) | 0.66 | 0.155 | 0.364 | 0.69 | 0.363 |
| Post-hoc t-test | (df*, σ) | (156, 0.12) | (92, 0.14) | (72, 0.12) | (71, 0.12) | (81, 0.14) |
| | p value | 0.235 | 0.108 | 0.352 | 0.183 | 0.732 |
| Proportion of cells with significant $r^2$ | pCA1 | 78/79 | 44/50 | 31/31 | 34/35 | 35/36 |
| | dCA1 | 78/79 | 40/44 | 43/43 | 36/38 | 47/47 |
| $\chi^2$ test for independence | $\chi^2$ | 0 | 0.208 | - | 0.267 | 1.32 |
| | p value | 1 | 0.648 | - | 0.604 | 0.25 |
| **TMIa for all place cells** | | | | | | |
| pCA1 median | | 0.86 | 0.87 | 0.82 | 0.83 | 0.87 |
| dCA1 median | | 0.86 | 0.82 | 0.86 | 0.84 | 0.84 |
| Median difference [95% CI] | | -0.0037 [-0.04, 0.03] | 0.05 [-0.012, 0.11] | 0.04 [-0.09, 0.04] | -0.017 [-0.06, 0.03] | 0.023 [-0.03, 0.09] |
| Wilcoxon rank sum test | z value | 0.163 | 0.973 | -0.219 | -0.215 | 1.3 |
| | rank sum | 6328 | 2504 | 1142 | 1275 | 1654 |
| | p value | 0.87 | 0.33 | 0.826 | 0.829 | 0.193 |
| Post-hoc Lilliefor's test | p value (pCA1) | 0.059 | 0.003 | 0.413 | 0.114 | 0.005 |
| | p value (dCA1) | 0.009 | 0.172 | 0.013 | 0.014 | 0.001 |
| Post-hoc t-test | (df, σ) | Failed test for normality | | | | |
| | p value | | | | | |
| Proportion of cells with significant TMIa | pCA1 | 78/79 | 46/50 | 30/31 | 31/35 | 36/36 |
| | dCA1 | 78/79 | 41/44 | 43/43 | 38/38 | 44/47 |
| $\chi^2$ test for independence | $\chi^2$ | 0 | 0.047 | 1.406 | 4.59 | 2.38 |
| | p value | 1 | 0.827 | 0.235 | 0.03 | 0.122 |

(*df = degrees of freedom, σ = standard deviation)
